# FAM35A co-operates with REV7 to coordinate DNA double-strand break repair pathway choice

**DOI:** 10.1101/365460

**Authors:** Steven Findlay, John Heath, Vincent M. Luo, Abba Malina, Théo Morin, Billel Djerir, Zhigang Li, Arash Samiei, Estelle Simo-Cheyou, Martin Karam, Halil Bagci, Dolev Rahat, Damien Grapton, Elise G. Lavoie, Christian Dove, Husam Khaled, Hellen Kuasne, Koren K. Mann, Kathleen Oros Klein, Celia M. Greenwood, Yuval Tabach, Morag Park, Jean-Francois Côté, Alexandre Maréchal, Alexandre Orthwein

**Affiliations:** Lady Davis Institute for Medical Research, Segal Cancer Centre, Jewish General Hospital, 3755 Chemin de la Côte-Sainte-Catherine, Montreal, Quebec, H3T 1E2, Canada; Division of Experimental Medicine, McGill University, Montreal, Quebec, H4A 3J1, Canada; Department of Microbiology and Immunology, McGill University, Montreal, Quebec, H3A 2B4, Canada; Gerald Bronfman Department of Oncology, McGill University, Montreal, Quebec, H4A 3T2, Canada; Department of Biology, Université de Sherbrooke, Sherbrooke, Quebec, J1K 2R1, Canada; Institut de Recherches Cliniques de Montréal (IRCM), Montreal, Quebec, H2W 1R7, Canada; Department of Developmental Biology and Cancer Research, The Institute for Medical Research Israel-Canada, The Hebrew University of Jerusalem, 91120, Jerusalem, Israel; Department of Epidemiology, Biostatistics and Occupational Health, McGill University, Montreal, Quebec, H3A 1A2, Canada; Rosalind and Morris Goodman Cancer Research Centre, McGill University, Montreal, Quebec, H3A 1A3, Canada; Department of Anatomy and Cell Biology, McGill University, Montreal, Quebec, H3A OC7, Canada; Département de Biochimie et Médecine Moléculaire, Université de Montréal, Montreal, Quebec, H3C 3J7, Canada; Département de Médecine (Programmes de Biologie Moléculaire), Université de Montréal, Montreal, Quebec, H3C 3J7, Canada

**Keywords:** DNA double-strand break, Non-Homologous End-Joining, DNA repair pathway choice, REV7

## Abstract

DNA double-strand breaks (DSBs) can be repaired by two major pathways: non-homologous end-joining (NHEJ) and homologous recombination (HR). DNA repair pathway choice is governed by the opposing activities of 53BP1, in complex with its effectors RIF1 and REV7, and BRCA1. However, it remains unknown how the 53BP1/RIF1/REV7 complex stimulates NHEJ and restricts HR to the S/G2 phases of the cell cycle. Using a mass spectrometry (MS)-based approach, we identify 11 high-confidence REV7 interactors and elucidate the role of a previously undescribed factor, FAM35A/SHDL2, as a novel effector of REV7 in the NHEJ pathway. FAM35A depletion impairs NHEJ-mediated DNA repair and compromises antibody diversification by class switch recombination (CSR) in B-cells. FAM35A accumulates at DSBs in a 53BP1-, RIF1- and REV7-dependent manner and antagonizes HR by limiting DNA end resection. In fact, FAM35A is part of a larger complex composed of REV7 and another previously uncharacterized protein, C20orf196/SHDL1, which promotes NHEJ and limits HR. Together, these results establish FAM35A as a novel effector of REV7 in controlling the decision-making process during DSB repair.

## INTRODUCTION

Due to their highly recombinogenic and pro-apoptotic potentials, DNA double-strand breaks (DSBs) are one of the most cytotoxic DNA lesions. Their inaccurate resolution can result in point mutations, small deletions/insertions, chromosomal rearrangements or loss of gross genetic information that drive genomic instability, carcinogenesis and cell death (reviewed in (Tubbs and Nussenzweig, 2017)). To avoid these deleterious outcomes, cells have deployed a complex network of proteins to signal and repair DSBs. One critical step during the DSB response consists in the choice between two mutually exclusive DNA repair pathways: Non-Homologous End Joining (NHEJ) and Homologous Recombination (HR) (reviewed in (Ceccaldi et al., 2016)). This decision process, named DNA repair pathway choice, integrates several elements including the cell cycle status, the complexity of the DNA end and the epigenetic context. Importantly, DNA repair pathway choice is under the control of two antagonizing factors, 53BP1 and BRCA1 (reviewed in (Hustedt and Durocher, 2016)).

NHEJ is predominantly involved in the repair of DSBs during the G1 phase of the cell cycle. It is characterized by a limited processing of the DNA ends catalyzed by the nuclease Artemis and their subsequent ligation by DNA ligase IV (reviewed in (Betermier et al., 2014)). Importantly, NHEJ is promoted by the recruitment of 53BP1 at DSBs, along with its effectors RIF1, REV7 and PTIP (Chapman et al., 2012; Callen et al., 2013; Chapman et al., 2013; Di Virgilio et al., 2013; Escribano-Diaz et al., 2013; Feng et al., 2013; Zimmermann et al., 2013; Boersma et al., 2015; Xu et al., 2015). These latter factors play a central in several additional biological processes, including the establishment of a protective immunity during class switch recombination (CSR), a programmed DSB-dependent process that specifically occurs in B-cells (Manis et al., 2004; Ward et al., 2004; Chapman et al., 2013; Di Virgilio et al., 2013; Escribano-Diaz et al., 2013; Boersma et al., 2015; Xu et al., 2015).

In S/G2 phases of the cell cycle (when sister chromatids are available as templates), HR is activated and can alternatively repair DSBs. One of the key features of HR is the formation of long stretches of single-stranded DNA (ssDNA), a process called DNA end resection (reviewed in (Fradet-Turcotte et al., 2016)). The resulting ssDNA stretches are rapidly coated by RPA, which is subsequently replaced by the recombinase RAD51 to form nucleofilaments that are a pre-requisite for the subsequent search of homology, strand invasion and strand exchange before the resolution of the DSB by the HR machinery. Critically, BRCA1 promotes the initiation of DNA end resection and HR-mediated DSB repair by preventing the recruitment of 53BP1 and its downstream effectors to sites of DNA damage in S/G2 phases (Chapman et al., 2012; Chapman et al., 2013; Escribano-Diaz et al., 2013; Feng et al., 2013), thereby antagonizing 53BP1 function in NHEJ.

While the opposing role of 53BP1 and BRCA1 in DNA repair pathway choice has been extensively scrutinized over the past years, it remains largely unclear how the 53BP1 downstream effectors, namely REV7, promote NHEJ and antagonize BRCA1-mediated HR in G1 phase of the cell cycle (Boersma et al., 2015; Xu et al., 2015). REV7 is an adaptor protein that has been described for its role in mitotic progression through the control of both the activity of the spindle assembly checkpoint (SAC) and the formation of a functional anaphase-promoting complex/cyclosome-Cdc20 (APC/C) (Cahill et al., 1999; Listovsky and Sale, 2013; Bhat et al., 2015). In parallel, REV7 is a well-defined player in DNA translesion synthesis (TLS) (reviewed in (Waters et al., 2009)) as well as DSB repair by HR as part of a complex composed of the deoxycytidyl (dCMP) transferase REV1 and the catalytic subunit of the DNA polymerase ζ, REV3L (Sharma et al., 2012). The recent discovery that REV7 participates in the NHEJ pathway in a TLS-independent manner raised fundamental questions about how this adaptor protein promotes DSB repair and controls DNA repair pathway choice.

In this present study, we sought to get insight into the decision-making process underpinning DNA repair pathway choice by deciphering the interactome of REV7. Using a mass spectrometry (MS)-based approach, we identified FAM35A/SHLD2 as a novel effector of REV7 in the NHEJ pathway. FAM35A accumulates at DSBs in a 53BP1-, RIF1- and REV7-dependent manner. Importantly, depletion of FAM35A impairs both NHEJ and CSR, while promoting DNA end resection and HR. In fact, FAM35A acts in concert with another previously uncharacterized protein C20orf196/SHLD1 in promoting NHEJ and antagonizing HR. Altogether, our results provide a new insight into the molecular events that control DNA repair pathway choice.

## RESULTS

### Mapping of REV7 proximal/interacting partners relevant for DNA repair pathway choice

To get better insight into the interactome of REV7, we performed a standard affinity purification (AP) followed by MS (AP-MS) (Fig EV1.A), where REV7 was tagged with the Flag epitope and stably expressed in the human embryonic kidney 293 (HEK293) cell line using the Flp-In/T-REX system (Fig EV1.B). As a complementary approach, we used a proximity-based biotin labelling technique (BioID), which allows the monitoring of proximal/transient interactions (Fig EV1.A) (Roux et al., 2012; Roux et al., 2013). Briefly, REV7 was fused to a mutant of an *E.coli* biotin-conjugating enzyme (BirA*) and stably expressed in HEK293 as previously described (Lambert et al., 2015). This fusion protein is capable of biotinylating proteins that come in close proximity or directly interact with REV7 (Fig EV1.C). Labelled proteins were subsequently purified by streptavidin affinity and identified by MS. Both approaches were carried out in triplicate using extracts of cells treated in the absence or presence of the radiomimetic DNA damaging drug Neocarzinostatin (NCS). We identified 140 high-confidence REV7 interactors that were either common to the four experimental conditions or found in both the AP-MS and the BioID following NCS treatment (Fig.1A and Table 1). As expected, pathways critical for mitosis and DNA repair were enriched in our list of REV7 partners (Fig EV1.D). To further refine REV7 interactors, we intersected our data with previously reported proteomic profiling of REV7 (Nelson et al., 1999; Chen and Fang, 2001; Weterman et al., 2001; Guo et al., 2003; Iwai et al., 2007; Zhang et al., 2007; Hong et al., 2009; Medendorp et al., 2009; Tissier et al., 2010; Vermeulen et al., 2010; Listovsky and Sale, 2013); Rolland et al. (2014); (Huttlin et al., 2015; Huttlin et al., 2017). Using this methodology, we obtained 11 high-confidence REV7 interactors (Fig.1B and Fig EV1.E), including the chromosome alignment-maintaining phosphoprotein (CHAMP1), a kinetochore-microtubule attachment protein that has been recently linked to REV7 and its role during mitotic progression (Itoh et al., 2011), as well as the cell-division cycle protein 20 (CDC20), a critical activator of the anaphase promoting complex (APC/C) that allows chromatid separation and entrance into anaphase (Chen and Fang, 2001; Listovsky and Sale, 2013; Bhat et al., 2015). However, whether these high-confidence REV7 interactors play any role in the DSB response remains unresolved.

**Figure 1.**
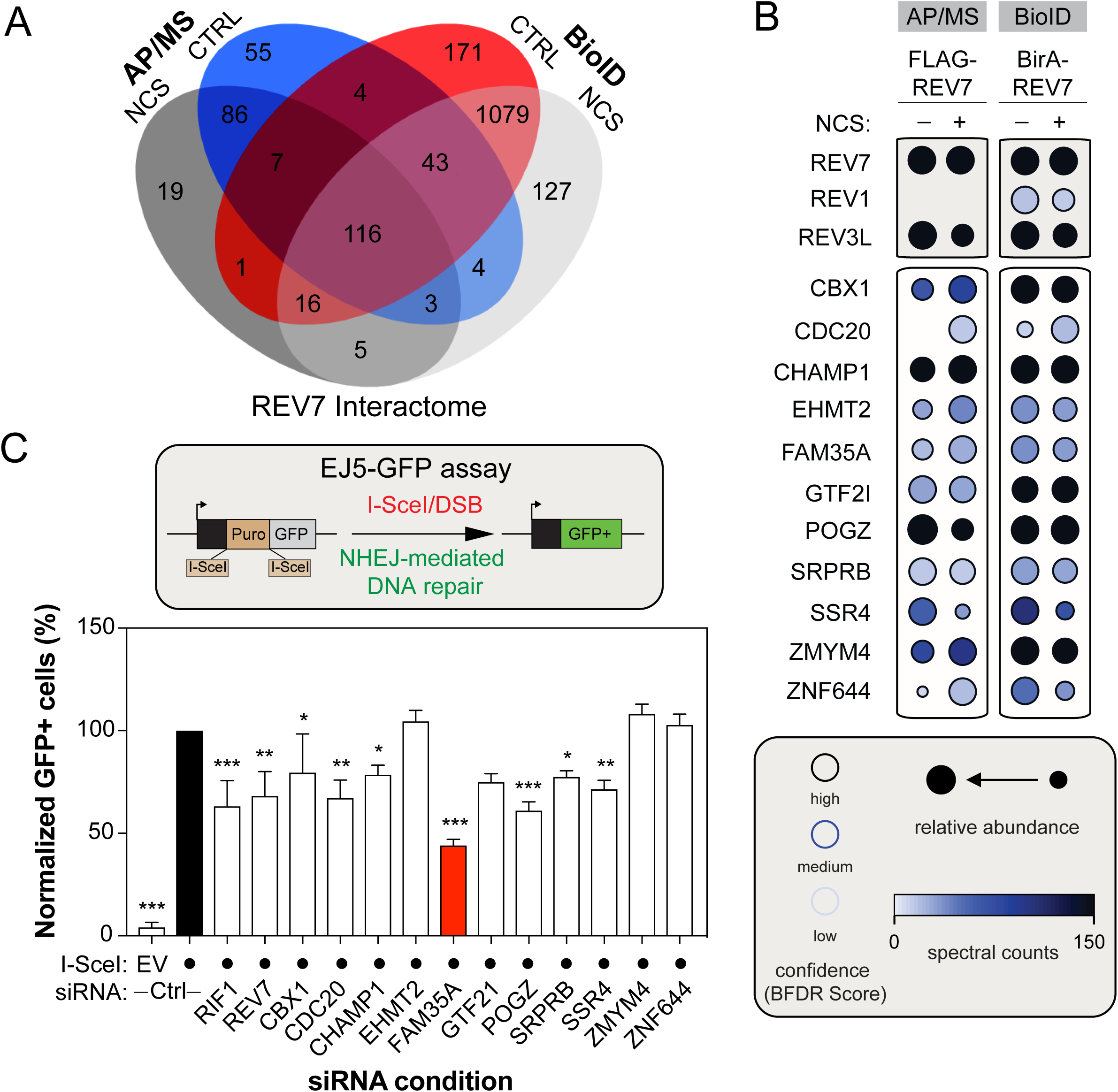
Identification of novel REV7 interactors relevant for the NHEJ pathway. (A) Venn Diagram representing the distribution of proteins identified by both the BioID and the standard AP/MS of REV7, with or without DNA damage (NCS). (B) Selected BioID REV7 results, shown as dot plots. The spectral counts for each indicated prey protein are shown as AvgSpec. Proteins were selected based on and iProphet probability of >0.95, BFDR of <0.05 and ≥10 peptide count. The circle size represents the relative abundance of preys over baits. (C) U2OS EJ5-GFP cells were transfected with the indicated siRNAs. At 24hr post-transfection, cells were transfected with the I-SceI expression plasmid, and the GFP^+^ population was analyzed 48 hr post-plasmid transfection. The percentage of GFP^+^ cells was determined for each individual condition and subsequently normalized to the non-targeting condition (siCTRL). Data is presented as the mean ± SD (n=3). Significance was determined by one-way ANOVA followed by a Dunnett test. *p<0.05, **p<0.005, ***p<0.0005

To ascertain the relevance of these interactors for NHEJ, we used a well-established GFP-based reporter assay that monitors total NHEJ events (Bennardo et al., 2008), the EJ5-GFP assay, and targeted each candidate using small interfering RNA (siRNA) pools (Fig.1C). As positive controls, we incorporated both RIF1 and REV7, which have been previously shown to impair this assay (Chapman et al., 2013; Escribano-Diaz et al., 2013; Boersma et al., 2015). Out of the 11 candidates tested, downregulation of seven REV7 interactors significantly impaired the restoration of the GFP signal following DSB induction and subsequent repair by NHEJ (Fig.1C), without impacting drastically cell cycle progression (Fig EV1.F). FAM35A emerged as our strongest hit, with a reduction of more than 60% of the GFP signal compared to the control condition in this assay (Fig.1C). Therefore, we concentrated our efforts on this factor to better define its involvement in DNA repair pathway choice.

### FAM35A promotes DSB repair in human cells

To get an evolutionary perspective and determine whether FAM35A may be relevant for DNA repair, we used a novel phylogenetic profiling (PP) approach and defined the landscape of genes that co-evolved with FAM35A among mammals and vertebrates (Tabach et al., 2013a). Importantly, this PP method has been previously shown to successfully predict protein function by analyzing the genes that co-evolved with a given factor of interest (Tabach et al., 2013a; Tabach et al., 2013b). Gene ontology analysis for biological process enrichment identified DNA repair, IL-1 signaling, Nucleotide Excision repair (NER) and the APC/C-CDC20 pathway as the most significant biological functions associated to genes that co-evolved with FAM35A (Fig.2A). Strikingly, FAM35A co-evolves with RIF1 in both mammalians and vertebrates, which further suggests a putative role of FAM35A in the maintenance of genome stability.

**Figure 2.**
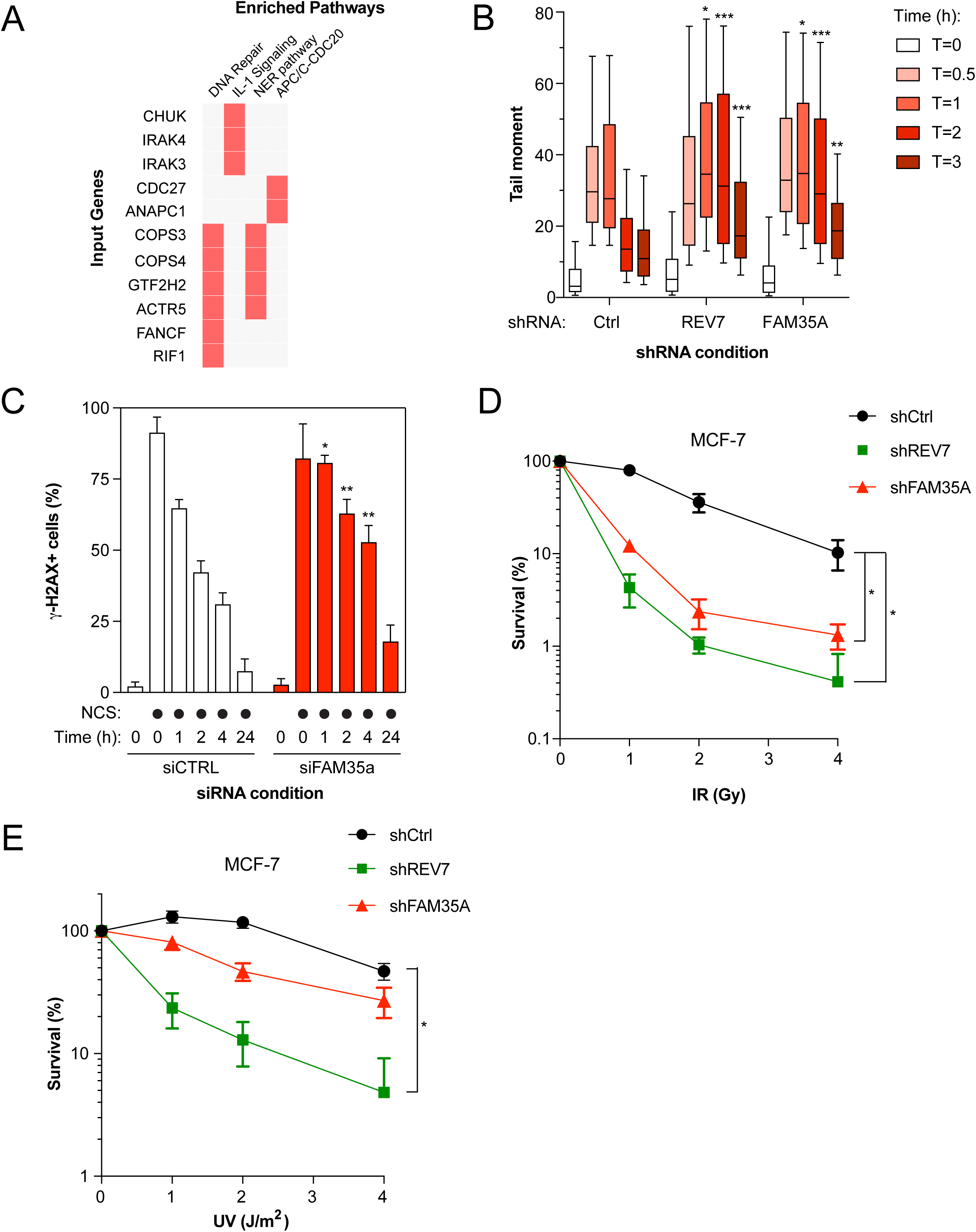
FAM35A plays a critical in DNA repair. (A) Pathway enrichment analysis on genes co-evolving with FAM35A in Mammalians and Vertebrates using a phylogenetic profiling approach followed by an Enrichr-based analysis. (B) Quantification of the Neutral Comet assay. U2OS cells stably expressing shCtrl, shREV7 or shFAM35A were exposed to IR (10 Gy) and run in low melting agarose under neutral conditions. Immunofluorescence against DNA stained with SYBR Gold was performed to measure the tail moment. Significance was determined by one-way ANOVA followed by a Dunnett test. *p<0.05, **p<0.005, ***p<0.0005. (C) U2OS cells were transfected with the indicated siRNA. 48h post-transfection the cells were treated with NCS to induce DNA damage and the cells were harvested at 0, 1, 2, 4 and 24 hr post NCS treatment. Flow Cytometry analysis of phosphorylated-H2AX signal was used to measure γ-H2AX endogenous signal. Significance was determined by two-way ANOVA followed by a Bonferroni test. *p<0.05, **p<0.005. (D) Sensitivity to IR monitored by Colony formation assay. MCF-7 cells stably expressing the the indicated shRNAs were exposed to increasing doses of IR. 24 h post-irradiation, the cells were re-seeded to allow colony formation, fixed and stained with 0.4% crystal violet. Shown is the quantification of colonies per condition which possessed more than 50 colonies. Significance was determined by one-way ANOVA followed by a Dunnett test. *p<0.05. (E) Similar to (D), except that cells were exposed to increasing doses of UV radiation. Significance was determined by one-way ANOVA followed by a Dunnett test. *p<0.05.

To explore this hypothesis, we first assessed the ability of FAM35A to promote DSB repair using the neutral comet assay. We observed that depletion of FAM35A by a short hairpin RNA (shRNA) in the osteosarcoma U2OS cell line resulted in the persistence of comet tails (time points 1, 2 and 3 hours) following exposure to ionizing radiation (IR; 10Gy), compared to control cells (Fig.2B and Fig EV2.A). For comparison, we used a shRNA targeting REV7 and obtained similar results. In a second assay, we monitored the phosphorylation of the histone variant H2AX (γ-H2AX), a marker of DSBs, over time by flow cytometry following treatment with NCS. Again, the kinetics of γ-H2AX resolution was delayed in FAM35A-depleted U2OS cells compared to control conditions (Fig.2C and Fig EV2.B), suggesting a role of FAM35A in DSB repair.

Next, we employed the breast cancer MCF-7 cell line to study the impact of FAM35A depletion on survival following DSB induction by IR. We observed that FAM35A depletion hypersensitizes MCF-7 cells to IR, in a manner similar to REV7 (Fig.2D). Loss or depletion of REV7 has been previously linked to a hypersensitivity to UV in line with its TLS function (Lawrence et al., 1985), raising the question of whether FAM35 is associated with a similar phenotype. Strikingly, FAM35A depletion did not sensitize MCF-7 cells to increased doses of UV (Fig.2E), suggesting that FAM35A is dispensable for TLS. From these results, we conclude that FAM35A is critical for DNA repair in a TLS-independent manner.

### FAM35A is recruited to DSBs through its N-terminal domain

FAM35A is a 904-amino-acid protein with very limited structural information available (Fig.3A). By performing structure prediction analyses on FAM35A protein sequence using Motif Scan (MyHits, SIB, Switzerland) and InterProScan5 (Jones et al., 2014), we identified a putative N-terminal DNA binding-domain (NUMOD3 domain) and a structural motif in its C-terminus that we labelled PFAM (Fig.3A). Recently, a structural prediction analysis of FAM35A also defined three putative OB-fold like domains at its C-terminus (Tomida et al., 2018). Finally, previous phospho-proteomic analysis identified a S/Q substrate of ATM/ATR following DNA damage at position 339 (Matsuoka et al., 2007).

**Figure 3.**
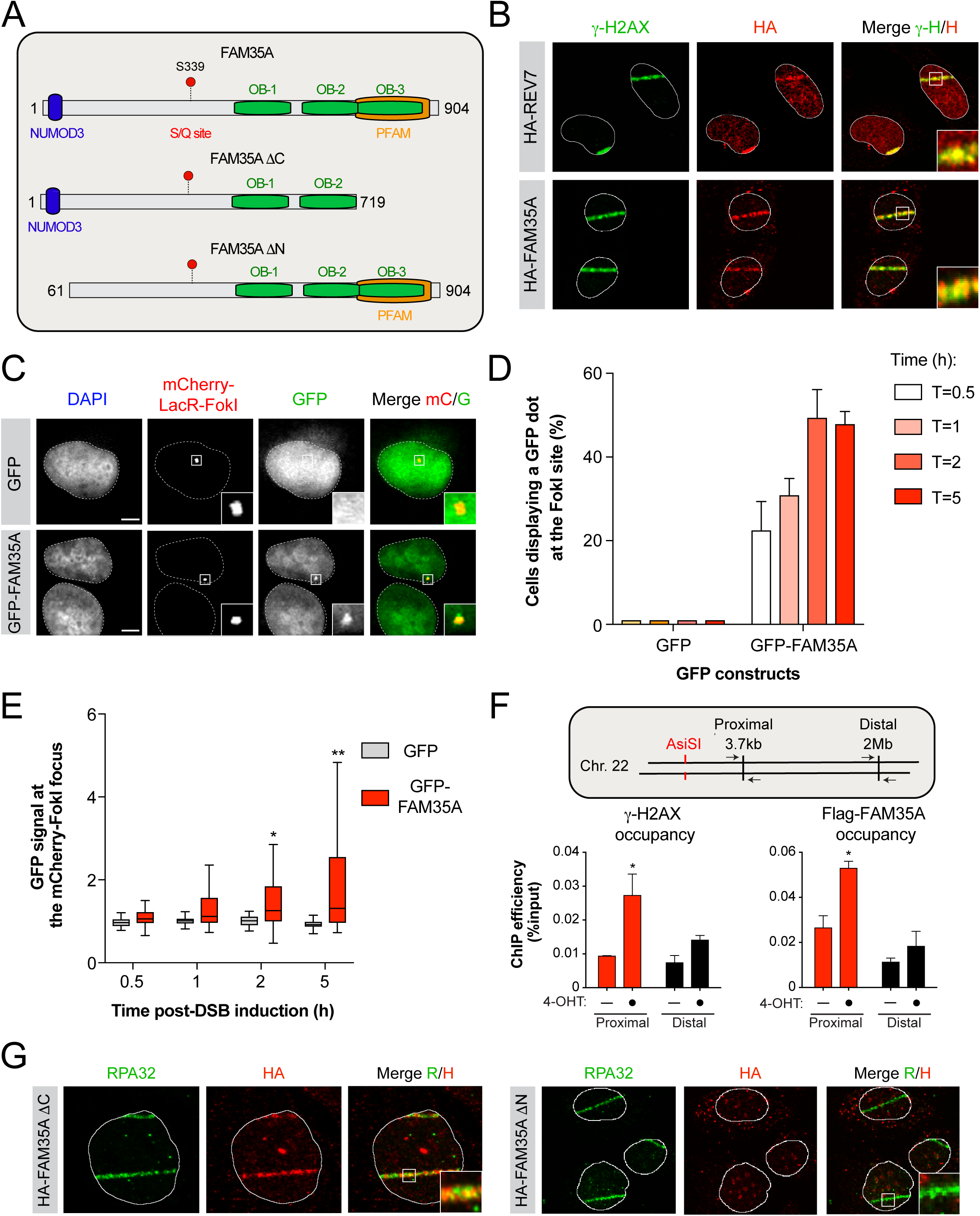
FAM35A is recruited and accumulates at DNA damage sites. (A) Schematic representation of FAM35A and its deletion mutants. (B) U2OS cells stably expressing HA-REV7 (Top) or HA-FAM35A (Bottom) were pre-sensitized with 10ug/mL Hoescht 33342 before exposed to UV micro-irradiation. Immunofluorescence against endogenous HA and γ-H2AX epitope was subsequently performed to monitor REV7 and FAM35A accumulation at sites of damage. Shown are representative micrographs. (C) U2OS LacR-Fok1 cells were transfected with GFP or GFP-FAM35A and 24 h later DNA damage was induced using Shield-1 and 4-OHT. The cells were then processed for GFP and mCherry immunofluorescence. Shown are representative micrographs. Scale bar = 5μm. (D) Quantification of the experiments shown in (C). Data are represented as the mean ± SD (n=3). At least 100 cells per condition were counted. (E) Quantification of the experiments shown in (C). Shown is the quantification of the GFP signal at the mCherry-LacR-Fok1 focus. Data are represented as a box-and-whisker plot in the style of Tukey. At least 100 cells per condition were counted. Significance was determined by non-parametric test followed by a Kruskal-Wallis test. *p<0.005, **p<0.0005. (F) Schematic representation of the site-directed generation of DSB by the restriction enzyme *AsiSI* (Top). 293T cell lines expressing ER-AsiSI with Flag-FAM35A and treated with 1 μM of 4- OHT. 6h later the cells were processed and immunoprecipitated with Anti-FLAG Magnetic Beads and anti-γ-H2AX.x/Protein A/G magnetic beads. DNA was purified and subjected to qPCR detection. Shown is the quantification of IP efficiency as the percentage of DNA precipitated from input (Bottom). Significance was determined by two-way ANOVA followed by a Sidak test. *p<0.05. (G) U2OS cells stably expressing HA-FAM35A-Δ C-Term (Left) or HA-FAM35A-Δ N-Term (Right) were processed as in (B). Immunofluorescence against endogenous HA and RPA32 epitope was subsequently performed to monitor RPA32 and FAM35A accumulation at sites of damage. Shown are representative micrographs.

To gain better insight into how FAM35A promotes DSB repair, we tested whether it accumulates at DNA damage sites. Indeed, we observed that HA-tagged FAM35A, similar to REV7, is rapidly recruited to laser microirradiation-induced DNA damage in U2OS cells, colocalizing with γ-H2AX (Fig.3B). As laser microirradiation elicits high levels of both single-strand breaks (SSBs) and DSBs, we complemented this approach by using the previously described mCherry-LacR-FokI-induced DSB reporter system (Tang et al., 2013). Here, GFP-tagged FAM35A is readily recruited to localized FokI-induced DSBs 30 min post-induction (Fig.3C) and the majority of cells displays a GFP-FAM35A-positive signal at the mCherry dot 2 hours post-DSB induction (Fig.3D). Furthermore, GFP-FAM35A accumulation at DSBs is significantly distinct from empty vector 2 hours following the induction of DNA damage (Fig.3E). We therefore used this experimental approach for our subsequent FokI-based experiments. As an orthogonal validation of the recruitment of FAM35A to DNA damage, we sought to use a well-established system where the induction of targeted DSBs is triggered by the controlled expression of the *Asi*SI restriction enzyme fused to a modified oestrogen receptor hormone-binding domain (Iacovoni et al., 2010). This model has been used to monitor the recruitment of DNA repair factors in the vicinity of DSBs and we confirmed by chromatin immunoprecipitation (ChIP) that induction of a DSB at chromosome 22 following addition of 4-hydroxy tamoxifen (4-OHT) triggers the formation of γ-H2AX proximally (3.7kb), but not distally (2Mb) from the site of damage (Fig.3F). Importantly, we observed that Flag-tagged FAM35A displays a similar distribution around the *Asi*SI-induced DSB. Altogether, these data indicate that FAM35A is persistently recruited at DSBs, confirming its role in DNA repair.

We further characterized the role of FAM35A during DSB repair by examining which domains of FAM35A are critical for its recruitment to DNA damage sites. We found that its C-terminal PFAM/OB3 domain and its S/Q motif (S339) are dispensable for FAM35A accumulation at DNA damage sites (Fig.3F, Fig EV3.A and B). However, deletion of the first 61 amino acids of FAM35A impairs its recruitment to DNA damage sites. Altogether, these data indicate that FAM35A accumulates at DSBs via a previously uncharacterized N-terminal NUMOD3 motif.

### FAM35A associates with REV7 to promote NHEJ and limit HR

To decipher the link between FAM35A and REV7, we tested the genetic requirements for the recruitment of FAM35A to DSBs using the FokI system. Depletion of 53BP1, RIF1 or REV7 by siRNA impaired its recruitment to a localized site of DNA damage (Fig.4A). However, we did not observe any impact on the recruitment of FAM35A to the FokI site following BRCA1 depletion (Fig.4A). These data indicate that FAM35A is acting downstream of REV7 in the NHEJ pathway.

**Figure 4.**
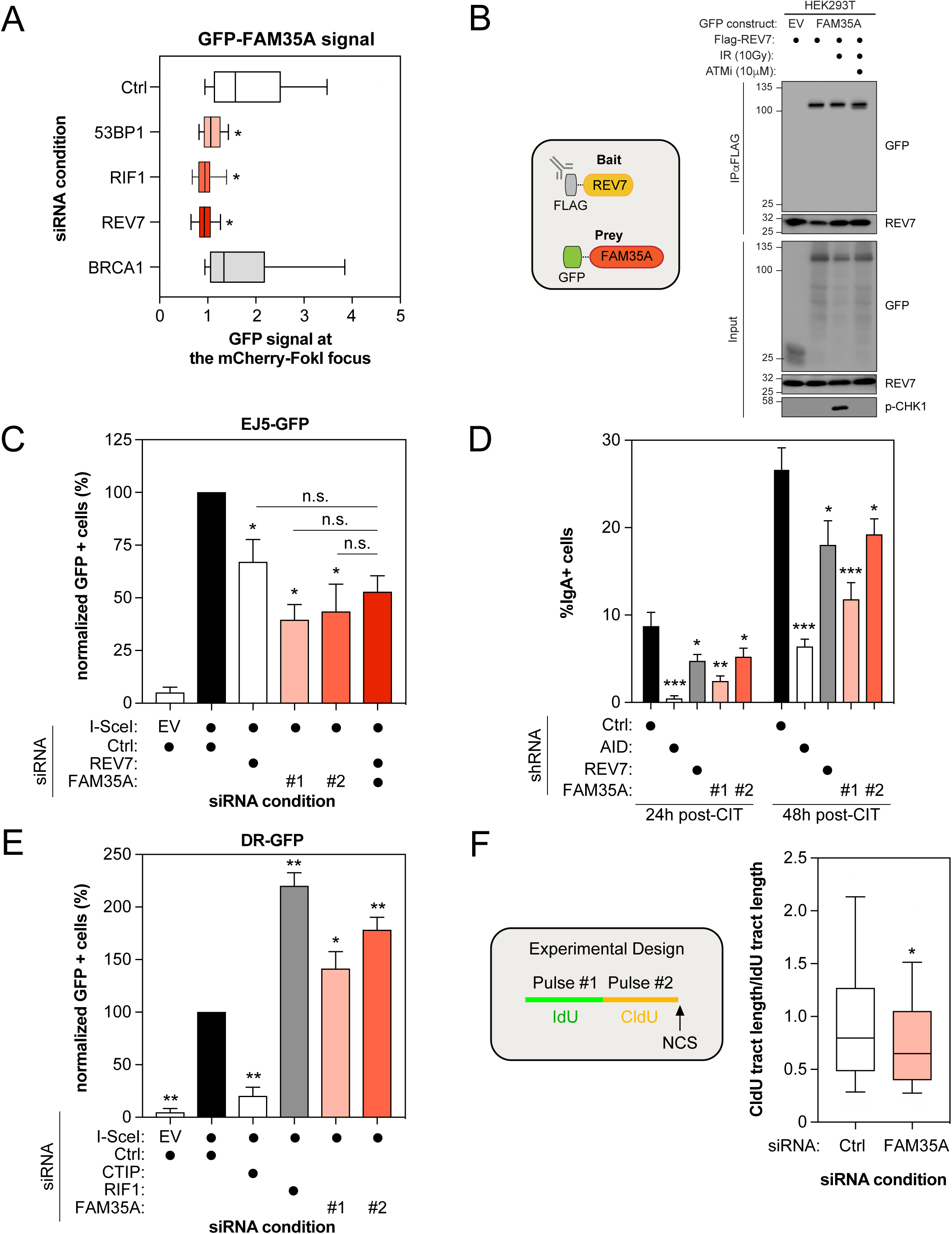
FAM35A is an effector of REV7 in promoting NHEJ and antagonizing HR. (A) U2OS mCherry-LacR-Fok1 cells were treated with the indicated siRNA and subsequently transfected with a GFP-FAM35A construct. 24 h post-transfection DNA damage was induced using Shield-1 and 4-OHT. The cells were then fixed and analyzed for the intensity of the GFP-FAM35A signal at mCherry-LacR-Fok1 focus. Shown is the quantification of the GFP-FAM35A signal at the Fok1 focus. Data are represented as a box-and-whisker plot where the whiskers represent the 10-90 percentile. At least 75 cells were counted per condition. Significance was determined by one-way ANOVA followed by a Dunnett test. *p<0.05. (B) 293T cells were transfected with Flag-REV7 and GFP-FAM35A expression vectors as indicated. 24h post-transfection cells were treated with DMSO or with 10uM of ATM inhibitor KU-60019 for 1 h prior to irradiation. 1h post-irradiation (10 Gy) nuclear extracts were prepared and REV7 complexes were immunoprecipitated using anti-Flag (M2) Resin and then analyzed by immunoblotting using GFP, REV7 and p-Chk1 antibodies. (C) U2OS EJ5-GFP cells were transfected with the indicated siRNAs. At 24hr post-transfection, cells were transfected with the I-SceI expression plasmid, and the GFP^+^ population was analyzed 48 hr post-plasmid transfection. The percentage of GFP^+^ cells was determined for each individual condition and subsequently normalized to the non-targeting condition (siCTRL). Data are presented as the mean ± SD (n=3). Significance was determined by one-way ANOVA followed by a Dunnett test using Ctrl+SceI as a comparison (*p<0.0005) or the indicated reference (n.s.= non-significant). (D) CH12F3-2 cells stably expressing the indicated shRNAs were stimulated with a cocktail of cytokines (CIT) to induce class switching to IgA. The percentage of IgA+ cells was monitored 24 and 48h post-stimulation by staining with an anti-IgA antibody followed by flow cytometry analysis. Data are presented as the mean ± SD (n=3). Significance was determined by one-way ANOVA followed by a Dunnett test. *p<0.05, **p<0.005, ***p<0.0005. (E) U2OS DR-GFP cells were transfected with the indicated siRNAs. At 24hr post-transfection, cells were transfected with the I-SceI expression plasmid, and the GFP^+^ population was analyzed 48 hr post-plasmid transfection. The percentage of GFP^+^ cells was determined for each individual condition and subsequently normalized to the non-targeting condition (siCTRL). Data are presented as the mean ± SD (n=3). Significance was determined by one-way ANOVA followed by a Dunnett test. *p<0.005, **p<0.0005. (F) Schematic representation of the DNA fiber assay experimental design (Left). U2OS cells were transfected with the indicated siRNAs and then treated with CldU, IdU and NCS 48h post-transfection as indicated. The slides were stained, dehydrated, mounted and visualized and shown is the quantification of CldU/IdU tract length in order to visualize DNA end-resection (Right). Data are represented as a box-and-whisker plot where the whiskers represent the 10-90 percentile. Significance was determined by one-way ANOVA followed by a Dunnett test. *p<0.0005.

We reasoned that if FAM35A is a direct effector of REV7, its recruitment to DSBs should be mediated through a physical interaction with REV7. Indeed, we confirmed the REV7-FAM35A interaction in co-immunoprecipitation experiments where tagged versions of both REV7 and FAM35A were expressed in 293T cells (Fig.4B). Exposure to IR did not stimulate the REV7-FAM35A interaction and pharmacological inhibition of ATM did not abrogate it (Fig.4B), suggesting that this interaction is constitutive and stable in 293T cells, which is consistent with our MS data.

Our data point towards a role of FAM35A in NHEJ downstream of REV7. Therefore, we confirmed that FAM35A depletion impairs NHEJ in the EJ5-GFP assay using two distinct siRNAs (Fig.4C). Next, we tested whether FAM35A and REV7 act epistatically to promote NHEJ. As expected, co-depletion of REV7 with FAM35A did not alter further the EJ5-GFP assay compared to the individual depletion (Fig.4C). We further defined the similarities between REV7 and FAM35A in the NHEJ pathway by testing the role of FAM35A in CSR. REV7 depletion has been previously shown to cause a profound defect in CSR in CH12F3-2 B-cells that switch from IgM to IgA following the addition of a cocktail of cytokines (IL-4/TGF-β/anti-CD40; CIT) and we confirmed these data (Fig.4D) (Boersma et al., 2015; Xu et al., 2015). Using two distinct shRNAs targeting FAM35A, we observed that its depletion impairs significantly CSR at both 24h and 48h post-activation (Fig.4D), suggesting that FAM35A regulates CSR at the level of DNA repair, which is consistent with its role as a REV7 effector.

53BP1 and its effectors have emerged as strong inhibitors of the HR pathway as well as the single-strand annealing (SSA) pathway (Chapman et al., 2012; Chapman et al., 2013; Di Virgilio et al., 2013; Escribano-Diaz et al., 2013; Feng et al., 2013; Zimmermann et al., 2013; Boersma et al., 2015; Xu et al., 2015). Therefore, we tested whether depletion of FAM35A alters both DNA repair pathways using the DR- and the SA-GFP reporter assays, respectively (Fig EV4.A) (Pierce et al., 1999; Stark et al., 2004). In both U2OS and HeLa DR-GFP cells (Fig.4E and Fig EV4.B), FAM35A depletion leads to a significant increase in HR using two distinct siRNAs, similar to what we observed with RIF1. Additionally, depletion of FAM35A, like RIF1, promotes SSA (Fig EV4.C) (Escribano-Diaz et al., 2013). This anti-HR role of 53BP1 and its effectors was attributed to a putative function in limiting DNA end resection, a key step in initiating DSB repair by HR. To define whether FAM35A controls DNA end resection, we carried out a modified version of the DNA combing assay, where a dual-pulse labelling of the replicating DNA was performed using two distinct nucleotides analogs (IdU and CldU) before addition of NCS (Fig.4F). While the length of the IdU-labeled DNA should not be altered by DNA end resection, we hypothesized that any increase in the processing of the DNA end should result in a shorter CldU-labeled DNA track and therefore a reduced ratio of CldU/IdU track length. Indeed, depletion of FAM35A in U2OS cells resulted in a significant reduction of the CldU/IdU ratio compared to control cells (Fig.4F and Fig EV4.D), strongly suggesting that FAM35A limits DNA end resection.

Loss of REV7 in BRCA1-deficient cells has also been shown to restore partially HR (Xu et al., 2015). We sought to examine whether depletion of FAM35A could result in a similar phenotype. Therefore, we co-depleted both BRCA1 and FAM35A in HeLa DR-GFP cells and we found a partial and significant restoration of HR in co-depleted vs. BRCA1-depleted cells (Fig EV4.E). Altogether, these results are consistent with a model where FAM35A, like 53BP1, RIF1 and REV7, promotes DSB repair by NHEJ and antagonizes HR by inhibiting DNA end resection.

### FAM35A associates with C20orf196 to promote NHEJ

It remains largely unclear how FAM35A promote NHEJ and limit HR, similar to 53BP1, RIF1 and REV7. Therefore, we determined the interactome of FAM35A using the BioID approach in presence or absence of DNA damage (+/-NCS; Fig.5A, Fig EV5.A and Table 2). Using this methodology, we identified previously described FAM35 interactors, including REV7, the RNA-binding protein HNRNPA1 and the E3 Ubiquitin ligase TRIM25 (Fig.5A) (Roy et al., 2014; Hein et al., 2015; Choudhury et al., 2017). Interestingly, several members of the COP9 signalosome (COPS4 and COPS6) and the Cullin-RING E3 Ubiquitin ligase family (CUL3, CUL4B, CUL5, DDB1) emerged as high-confidence proximal interactors of FAM35A. However, by comparing both REV7 and FAM35A BioID datasets, we did not identify any common complex of relevance for DNA repair. Therefore, we sought to undertake a more systematical and unbiased approach to identify novel DNA repair factors using the CRISPR/Cas9 technology (Fig.5B). We employed the previously described TKO.v1 sgRNA library that contains 91,320 sequences and targets 17,232 genes and applied it to an hTERT immortalized retinal pigment epithelial RPE1 cell line stably expressing Cas9 (Hart et al., 2015). To identify genes that are relevant for DNA repair, we used the chemotherapy drug doxorubicin as a selective agent (Fig.5B). TP53, along with CHK1 and TOP2A emerged as our strongest hits providing resistance to doxorubicin (Fig.5C). Their depletion was previously shown to elicit doxorubicin resistance (Burgess et al., 2008), thereby validating our approach (Fig.5C). We subsequently focused our analysis on doxorubicin-sensitizers, as they are likely to play a key role in DNA repair. Interestingly, C20orf196/SHLD1 scored as one of our most depleted genes (Fig.5C). This factor is of particular interest as it has been previously identified in the proteomic analysis of two of our high-confidence FAM35A interactors, REV7 (Hutchins et al., 2010) and CUL3 (Bennett et al., 2010). We therefore concentrated our efforts on this factor to define its link with FAM35A during DNA repair.

**Figure 5.**
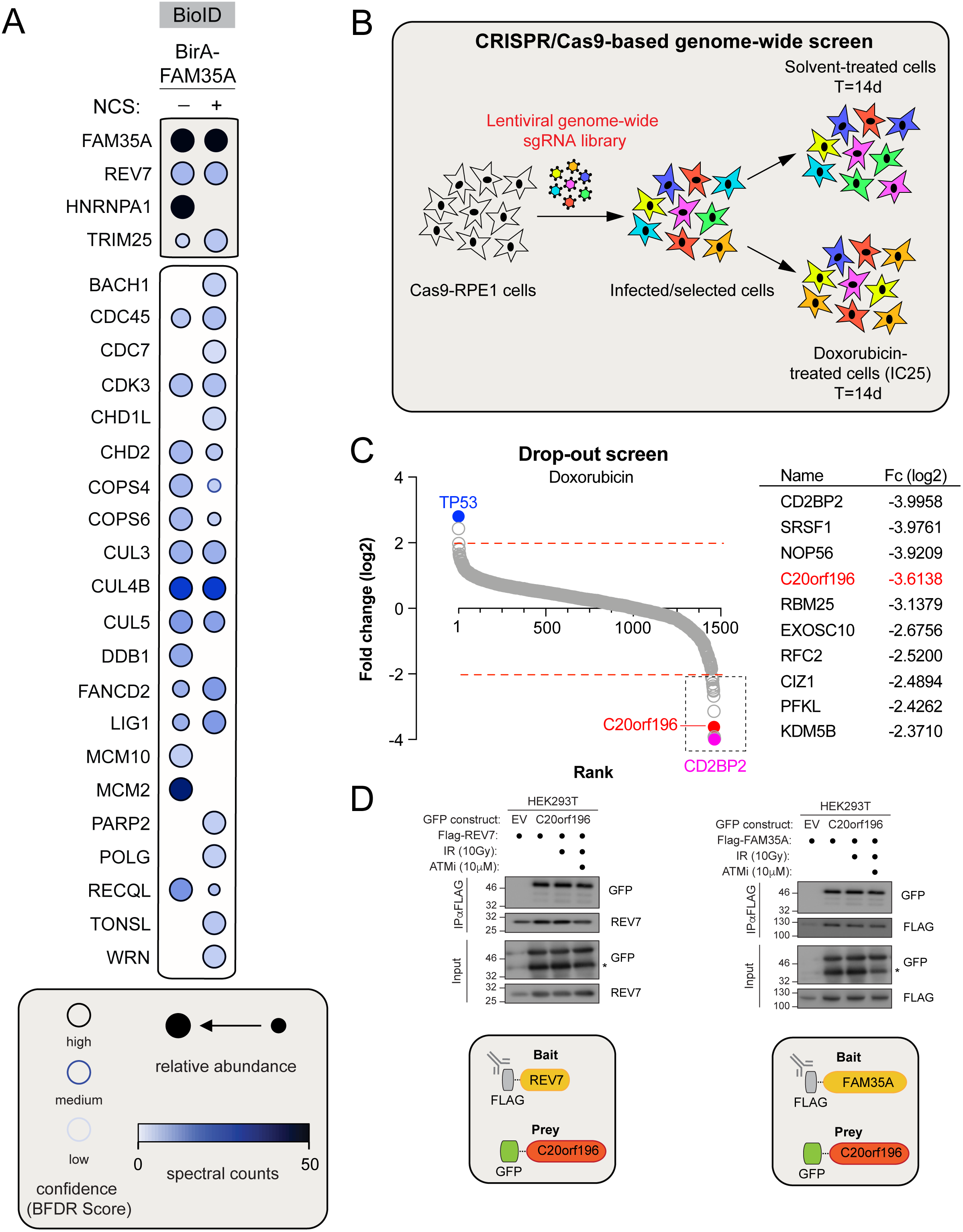
C20orf196 cooperates with FAM35A and REV7 to promote NHEJ and restrict HR. (A) Selected BioID results of FAM35A, shown as dot plots. The spectral counts for each indicated prey protein are shown as AvgSpec. Proteins were selected based on and iProphet probability of >0.95, BFDR of <0.05 and ≥10 peptide count. The circle size represents the relative abundance of preys over baits. (B) Schematic Representation of CRISPR/Cas9-based Genome-Wide Screen under Doxorubicin treatment. (C) Genes significantly enriched or dropped out after a 14-day treatment with Doxorubicin were identified by plotting as a Log2 fold change compared to untreated. Ranking was determined based on the Log2 fold score (Left). The top ten Doxorubicin sensitizers are indicated on the right with their respective fold change (Fc) in Log2. (D) 293T cells were transfected with Flag-REV7 and GFP-C20orf196 (Left) or Flag-FAM35A and GFP-C20orf196 (Right) expression vectors as indicated. 24h post-transfection cells were treated with DMSO or with 10uM of ATM inhibitor KU-60019 for 1 h prior to irradiation. 1h post-irradiation (10 Gy) nuclear extracts were prepared and REV7 or FAM35A complexes were immunoprecipitated using anti-Flag (M2) Resin and then analyzed by immunoblotting using GFP and REV7 antibodies.

First, we confirmed that REV7 and C20orf196 interact together by co-immunoprecipitation experiments where tagged versions of both REV7 and C20orf196 were expressed in 293T cells (Fig.5D). Next, we tested whether FAM35A interacts with C20orf196 using a similar approach (Fig.5D). Importantly, both REV7-C20orf196 and FAM35A-C20orf196 interactions did not increase upon IR treatment, neither did the pharmacological inhibition of ATM abrogates them (Fig.5D), similar to what we observed previously with the REV7-FAM35A interaction. If C20orf196 is part of a complex composed of REV7 and FAM35A, we would expect C20orf196 to accumulate at sites of damages, recapitulating the observations we made with both REV7 and FAM35. Therefore, we carried out laser stripe micro-irradiation experiments and observed that HA-tagged C20orf196 colocalizes with γ-H2AX at DNA damages sites (Fig EV5.B). We further investigated the role of C20orf196 in DNA repair using the EJ5- and the DR-GFP reporter assays. Depletion of C20orf196 resulted in a significant reduction of DSB repair by NHEJ in the EJ5-GFP assay (Fig EV5.C), as previously observed with FAM35A and REV7. Concomitantly, depletion of C20orf196 led to a significant increase in HR in the DR-GFP assay (Fig EV5.D). Altogether, these data suggest that FAM35 is a part of large multiprotein complex, composed of at least REV7 and C20orf196, that promotes NHEJ and restricts HR.

### FAM35A levels correlate with a poorer prognosis in a subset of breast cancer patients

Dysregulation of DSB repair pathways has been frequently observed in several types of cancer and extensively documented for its role in the pathobiology of breast cancer (BC). We sought to determine whether FAM35A may contribute to the outcome of BC by interrogating two distinct patient-based cohorts of triple negative breast cancer (TNBC) and basal-like BC.

Interestingly, high levels of FAM35A correlates with a poorer survival probability in a well annotated cohort of 24 TNBC patients (Fig.6A). We confirmed this observation in the publicly available TCGA database where we focused our analysis of basal-like BC patients. Again, high expressers of FAM35A have significantly lower relapse-free survival in this cohort (Fig.6B), suggesting a putative role of FAM35A in the pathobiology of a BC subset.

**Figure 6.**
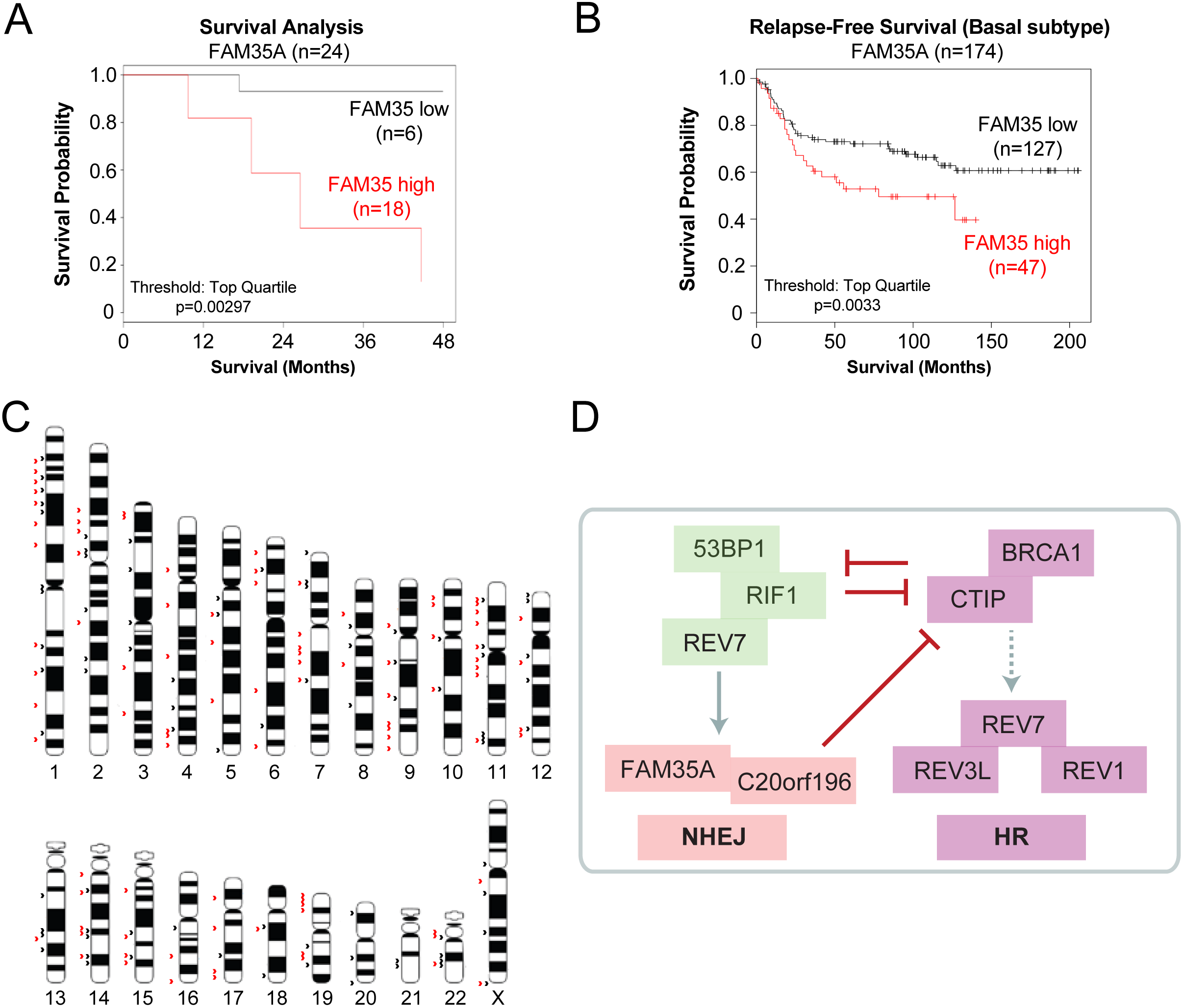
Relevance of FAM35A in breast cancer and HR restoration. (A) Survival analysis of low and high expressers of FAM35A in a cohort of 24 patients affected by Triple Negative Breast cancer (TNBC). Data are represented as Kaplan-Meyer curves with expression classified as low and high. Threshold for high cut-off is the top quartile. Significance was determined by calculating the hazard ratio with 95% confidence and the logrank P value. (B) Relapse-free Survival (Basal Subtype) of low and high expressers of FAM35A obtained from the KM-plotter database. Data are represented as Kaplan-Meyer curves with expression classified as low and high. Threshold for high cut-off is the top quartile. Significance was determined by calculating the hazard ratio with 95% confidence and the logrank P value. (C) Schematic representation of the top FAM35A peaks obtained by ChIP-seq. Black arrows indicate the location of the peaks obtained in normal conditions, while red arrows are representing the location of the FAM35A peaks detected following exposure to NCS. (D) Schematic incorporating FAM35A and C20orf196 as REV7 effectors in the NHEJ pathway and modulators of DNA repair pathway choice.

To better understand how high levels of FAM35A may promote BC carcinogenesis, we sought to examine the genome-wide occupancy of FAM35 by ChIP sequencing (ChIP-seq). We over-expressed a Flag-tagged version of FAM35A in HEK293 cells treated or not with NCS and used the Flag empty vector as background control. With this approach, we identified 441 distinct FAM35A peaks in our untreated conditions, while exposure to NCS resulted in the formation of 704 distinct peaks (Table 3). By mapped them across the genome, we observed very limited overlap between our two experimental conditions (58 overlapping binding sites; Fig.6C and Table 3), which confirmed that FAM35A re-localizes throughout the genome upon induction of DNA damage. Next, we tested whether FAM35A occupancy on chromatin requires a specific DNA motif. Therefore, we performed a motif analysis of our ChIP-seq data using the HOMER software and identified multiple binding motifs with high confidence in both experimental conditions (Table 4), suggesting that FAM35A binds DNA in a sequence-independent manner. Interestingly, CTCF-binding motifs are highly enriched in normal conditions, which may reflect the occupancy of FAM35A at common fragile sites (Canela et al., 2017). Indeed, it has been reported that chromosomal loops bound by both CTCF and Cohesin are highly vulnerable to the formation of DSBs (Canela et al., 2017) and FAM35 accumulation at these DSBs could drive genomic instability and carcinogenesis. Altogether, our data are consistent with a model where FAM35A needs to be tightly regulated to control DNA repair pathway choice where it acts as a downstream effector of REV7 in the NHEJ pathway and restricts DNA end resection, thereby antagonizing HR (Fig.6D).

## DISCUSSION

Two main DNA repair pathways, NHEJ and HR, are typically mobilized to repair cytotoxic DSBs and optimal pathway selection is central in preserving genome integrity. Several factors including 53BP1, RIF1 and REV7, emerged recently as key players in DNA repair pathway choice (Chapman et al., 2012; Chapman et al., 2013; Di Virgilio et al., 2013; Escribano-Diaz et al., 2013; Feng et al., 2013; Zimmermann et al., 2013; Boersma et al., 2015; Xu et al., 2015). However, it remains largely unclear how they modulate the proper balance between NHEJ and HR. In this study, we provide evidence of a novel complex composed of REV7 and two previously uncharacterized factors, FAM35A and C20orf196, which operates at DSBs to promote NHEJ while limiting DNA end resection, thereby antagonizing HR.

First, we identify FAM35A as an effector of REV7 in the NHEJ pathway. An unbiased co-evolutionary analysis (Tabach et al., 2013a; Tabach et al., 2013b) of FAM35 across 100 vertebrate species reveal a distinct pattern that is similar to other NHEJ genes, including RIF1, which indicates its association with DNA repair mechanisms. Through its N-terminal domain, FAM35A is mobilized to and accumulates at sites of DNA damage in a 53BP1-, RIF1- and REV7-dependent manner. Moreover, in a series of functional studies, we show that FAM35A is critical during both antibody diversification and DSB repair by the NHEJ pathway. Importantly, FAM35A and REV7 act together in an epistatic manner, which strongly indicates that FAM35A is promoting classical NHEJ downstream of REV7 (Fig.6D). We further show that, similar to 53BP1 and RIF1 (Chapman et al., 2012; Chapman et al., 2013; Di Virgilio et al., 2013; Escribano-Diaz et al., 2013; Feng et al., 2013; Zimmermann et al., 2013), FAM35A opposes HR by limiting DNA end resection. However, whether this anti-HR function of FAM35A is related to a steric hindrance of the DNA ends or an active process of preventing CtIP and its associated nucleases to initiate DNA end resection remains an avenue of investigation.

Second, our genome-wide screening approach identifies novel players in DNA repair, including C20orf196/SHLD1. Our data point toward a more complex model where REV7, FAM35A and C20orf196 cooperate together to promote NHEJ and limit HR. Indeed, we show that C20orf196 co-immunoprecipitates with both REV7 as previously described (Hutchins et al., 2010) but also FAM35A. Surprisingly though, we did not identify C20orf196 as a high confidence interactor of FAM35A in our BioID approach, likely due to the low abundance of this factor and in accordance with a previous report (Gupta et al., 2018). Still, our genetic dissection of C20orf196 recapitulates the striking data that we observe with FAM35A: (i) C20orf196 is recruited to and accumulates at sites of DNA damage; (ii) its depletion impairs NHEJ while promoting HR in two distinct GFP-based reporter assays. Why REV7 requires several factors to promote NHEJ and inhibit DNA end resection is unclear; our model suggests that, alike the Shelterin complex at telomeres (Schmutz and de Lange, 2016), which lacks any catalytic activity *per se*, REV7 forms a large multi-protein complex at DSBs to protect DNA ends from extensive processing and promote their rapid joining by the NHEJ machinery.

Finally, our observation that FAM35A levels correlate with a poor prognosis in a subset of BC has profound implications for the diagnosis and treatment of these patients. Imbalance in DSB repair pathways has been well documented to predispose and promote the development of BC; in the majority of the cases, inactivation of HR factors is the cause of this predisposition with a very limited understanding of the molecular mechanisms underlining this phenomenon. Our study points toward an expressional dysregulation of FAM35A as a potential predisposing factor to TNBC/Basal-like BC outcome, which may point toward a direct contribution of this novel NHEJ component in the pathobiology of BC. It will be of great importance to further define the role of FAM35A in BC as it may be a relevant biomarker for its diagnosis. Our ChIP-seq data suggest that FAM35A accumulates at CTCF-binding sites and could thereby drive genomic instability by promoting chromosomal translocations at break point cluster regions ((Canela et al., 2017).

Altogether, the work presented here not only uncovers two novel DNA repair factors that cooperate with REV7 in promoting NHEJ and antagonizing HR but it also provides the first evidence that FAM35A could benefit clinicians as a relevant biomarker for a subset of BC. Our data points toward a more complex model for DNA repair pathway choice where REV7 mobilizes additional factors to DSBs to catalyze NHEJ and limit the processing of DNA ends, thereby restricting HR to the S/G2 phases of the cell cycle.

## MATERIALS AND METHODS

### Cell culture and plasmid transfection

HEK293-T, Flp-In T-REx, -XT, and HeLa cells were cultured in Dulbecco’s Modified Eagle medium (DMEM; Wisent) and were supplemented with 10 % fetal bovine serum (FBS) and 1% Penicillin-Streptavidin (P/S). CH12F3-2 cells were cultured in RPMI 1640 (Wisent) supplemented with 10% FBS, 5% NCTC-109 media (Thermo Fisher), 50μM 2-mercaptoethanol and 1% P/S. U2OS cells were cultured in McCoy’s 5A Modified medium (Wisent) and was supplemented with 10 % FSB and 1% P/S. All cell lines were tested for mycoplasma contamination and STR DNA authenticated. Plasmid transfections were carried out using Lipofectamine 2000 Transfection Reagent (Invitrogen) following the manufacturer’s protocol. Lentiviral infections were done as previously described (Escribano-Diaz et al., 2013), with modifications listed below.

To generate the ER-AsiSI-expressing HEK293T cell line, retroviral particles were produced using pBABE HA-AsiSI-ER (kind gift of Dr. Michael Witcher, McGill University), the packaging plasmid pUMVC (addgene 8449), and the envelope plasmid VSV-g (Addgene #8454) co-transfected into HEK293T cells. To produce U2OS stable cell lines for microirradiation, lentiviral particles were produced by polyethyleneimine-mediated transfection of pHAGE-EF1α plasmid-cDNA with psPAX2 and pMD2.G packaging and envelope plasmids in 293 XT packaging cells. U2OS cells were infected with lentiviruses for 24h in media containing 8 μg/mL polybrene and 1μg/mL puromycin selection was applied. To generate a HEK293-TREx Flp-in cells were co-transfected with pOG44 and pcDNA5 FRT/TO FLAG-FAM35A and selected in with 200 μg/ml hygromycin B and 5 μg/ml blasticidin. The U2OS-LacI-FokI-mCherry cell line was a kind gift of R. Greenberg (University of Pennsylvania). The DNA-repair reporter cell lines DR-GFP, EJ5-GFP, and SA-GFP were a kind gift of Dr. Jeremy Stark (City of Hope National Medical Center, California).

### Plasmids

The cDNAs of human FAM35A, C20ORF196 and REV7 were obtained from Sidong Huang (McGill University). Quikchange site directed mutagenesis (Agilent) was performed as per manufacturers guidelines to obtain the FAM35A ΔC, FAM35A S339A, FAM35A ΔΝ mutants. All these constructs were transferred from ENTRY vectors into lentiviral pHAGE-EF1α vectors in frame with N-terminal 3XHA epitopes, GFP-construct, BirA-construct and Flag-tagged constructs using LR Clonase II according to manufacturer’s instructions (ThermoFisher). Plasmids encoding, I-SceI or pDEST-FRT-FLAG (EV) for the different GFP reporter assays, were kindly provided by Dr. Daniel Durocher (Lunenfeld-Tanenbaum Research Institute). The following pLKO-puro shRNA lentiviruses obtained from Mission library clones (Sigma) against mouse genes: Negative control (scramble); *Mad2l2* 45 (TRCN0000012845); *Aicda* (TRCN00000112033); *Fam35a* (TRCN0000183111); and *Fam35a* (TRCN0000183379).

### RNA interference

All siRNAs employed in this study were single duplex siRNAs purchased from Dharmacon (GE Healthcare, Colorado, US). RNAi transfections were performed using Lipofectamine RNAiMax (Invitrogen) in a forward transfection mode. Except when stated otherwise, siRNAs were transfected 48 h prior to cell processing. The individual siRNA duplexes used are: Control (D-001810-03); RIF1 (D-027983-02); CBX1 (L-009716-00); CDC20 (L-021601-02); CHAMP1 (L-021601-02); EHMT2 (L-006937-00); GTF21 (L-013686-00); POGZ (J-006953-10/12); REV1 (D-008234-01/02/0/3/04); REV3L (D-006302-01/02/03/04); REV7 (J-003272-14); SRPRB (L-013646-00); SSR4 (L-012264-00); ZMFM4 (L-019932-02); ZNF644 (L-007085-02); FAM35A (D-013761-01/02/03/04); C20orf196 (D-018767-01/02/03/04); BRCA1 (D-003461-05); CTIP (M-011376-00). In most of the experiments, FAM35A siRNAs D-013761-01and D-013761-03 were used except during the validation screen.

### Immunofluorescence microscopy

In most cases, cells were grown on glass coverslips. All steps were carried out at room temperature. Cells grown on coverslips were fixed in freshly prepared 2% paraformaldehyde for 10 minutes. Fixed cells were then incubated for 10 minutes with a combination of permeabilization/blocking buffer [0.1% Triton X-100 and 1% bovine serum albumin (BSA]. Next, primary antibodies were added for 1.5 hours in phosphate-buffered saline (PBS) + 1% BSA followed by three washes with PBS. Secondary antibody was next added in the same buffer for a period of one hour. Nuclei were stained with DAPI (1 μg/ml) for 5 min and subjected to a set of final washes with PBS and subsequently sterile water. After this, coverslips were mounted onto glass slides using a ProLong Diamond antifade reagent (Life Technologies). Images were acquired using a Zeiss LSM800 confocal microscope. Images were analyzed and quantified using ImageJ software [National Institutes of Health (NIH)]. For the FOK1 system, DSBs were induced by adding Shield-1 and 4-OHT for two hours prior to immunofluorescence sample preparation.

### Clonogenic Assay

Clonogenic assays was performed as described (Orthwein et al., 2014). Briefly, cells were allowed to reach ~50% confluence prior to genotoxic insult. Culture plates were then exposed to the indicated dose of IR or 254 nM ultraviolet (UV) light and allowed to recover overnight. Cells were then trypsinized and re-seeded into 60 cm dishes at 400 cells (or 800 cells for the highest dose) per dish. Colonies were allowed to form over the duration of 2 weeks and then fixed in 100% methanol and stained with 0.4% crystal violet (in 20% methanol). Colony number was manually tabulated with only colonies of >50 cells included in the total count.

### Co-Immunoprecipitation

HEK293T cells were co-transfected with pDEST-FRT-TO-Flag and pDEST-FRT-TO-GFP tagged vectors. Twenty-four hour post-transfection all cells were exposed to irradiation (10 Gy). ATMi-treated cells were exposed to 10 μM KU-60019 one hour prior to irradiation. Cells recovered at 37°C for one hour before being harvested and lysed in a high salt lysis buffer (50 mM Tris, 300 mM NaCl, 1mM EDTA, 1% Triton), supplemented with 1x Protease Inhibitor Cocktail (Roche)/Phosphate Inhibitor cocktail (Sigma) and gently rotated for 30 minutes at 4°C. Nuclear fractions were extracted with 0.25M CaCl_2_ and 250U benzonase and homogenized on an orbital shaker for 15 minutes at 30°C. The resulting solution was pelleted at 4°C at 18000g for 15 minutes and the supernatant was applied to an anti-Flag (M2) resin (Sigma) and equilibrated at 4°C for two hours. The anti-Flag resin was then washed once with the high salt lysis buffer and twice with the immunoprecipitation (IP) buffer (50mM Tris, pH 7.6, 150mM NaCl, 1mM EDTA). The immunoprecipitated proteins were eluted from the resin with 1x LDS NuPage sample buffer (10mM Tris-HCl, 140 mM Tris-base, 0.5 mM EDTA, 1% lithium dodecyl sulfate, 10% glycerol).

### Biotin labelling and sample preparation for MS

Samples for BioID were processed as previously described (Lambert et al., 2015). Briefly, HEK293-T cells were either transiently transfected with FLAG-BirA*-FAM35A or stably expressed using the T-REX system (FLAG-BirA*-REV7). Media was supplemented, 24 hours post-transfection, with 50μM biotin and cells were incubated for an additional 24 hrs with neocarzinostatin (NCS, 150 ng/ml). Cells were then harvested, washed twice with PBS, and dried. Pellets were subsequently resuspended in cold RIPA buffer containing: 50 mM Tris-HCl pH 7.4, 150 mM NaCl, 1 mM EDTA, 1% NP-40, 0.1% SDS, 0.5% sodium deoxycholate, 1mM PMSF, 1mM dithiothreitol, 1:500 Sigma-Aldrich protease inhibitor cocktail P8340. Cell homogenates were sonicated, followed by the addition of 250 U benzonase and centrifuged (12,000 g, 30 min). Supernatants were incubated with pre-washed Streptavidin-sepharose beads (GE, #17-5113-01) at 4°C with rotation for three hours. Beads were collected by centrifugation (2,000 rpm, 1 min), washed twice with RIPA buffer, three times with 50 mM ammonium bicarbonate (ABC, pH 8.2). Beads were resuspended in 50 mM ABC and treated with 1 μg trypsin (Sigma-Aldrich T6567) overnight at 37 °C with rotation. Digestion was continued by adding an additional 1 ug of trypsin for an additional 2 hrs at 37 °C with rotation. Supernatant containing peptides, and supernatants from two following washes with HPLC-grade H2O, was collected and pooled. Digestion was ended with the addition of formic acid to a final concentration of 5 %. Samples were centrifuged (13200rpm for 10 min) and the supernatants were dried in a SpeedVac for three hours at high rate. Peptides were resuspended in 5 % formic acid and kept at −80 °C for mass spectrometric analysis. MS processing and protein analysis were carried out as previously described.

Mass spectrometry data generated by the Regional Mass Spectrometry Centre (Université de Montréal) or the IRCM Proteomics Discovery Platform were stored, accessed, searched and analyzed using the ProHits laboratory information management system (LIMS) platform. Significance Analysis of INTeractome (SAINT)express (v3.6.1) was the statistical tool utilized to calculate the probability of protein-protein interaction from background, non-specific interactions (Choi et al., 2011). These results were evaluated with the Trans-Proteomic Pipeline (TPP v5.1) via the iProphet search engine integrated in the ProHits software (Liu et al., 2010; Shteynberg et al., 2011). A minimum of two unique peptide ions, an iProphet probability of >0.95, a Bait False Discovery Rate (BFDR) of <0.05, and a ≥10 peptide count were the criteria required for protein consideration. Biofilters were applied against albumin, artifact protein, cytoskeleton, and keratin. Resultant proteins from AP-MS (FLAG-REV7) and BioID (BirA-REV7; BirA-FAM35A) experiments were tabulated and analyzed for common potential interactors between the AP-MS and BioID groups, respectively. Common candidates were then sorted according to largest peptide counts. This analysis yielded 140 mutual candidates for the NCS+ group and 170 for the NCS-group. By way of literature review and the use of the Biological General Repository for Interaction Datasets (BioGRID), promising candidates were selected for targeted experimentation. Selected prey proteins were used for dot-plot heat map generation (see Figure 1 and 5). Plots were generated using the ProHits Visualization Suite (ProHits-viz) (Knight et al., 2017).

### GFP-based DNA repair assays

For DR-, EJ5-, EJ2-, SA-GFP reporter assays, U2OS or HeLa cells carrying the respective GFP expression cassette were transfected with indicated siRNA. Twenty-four hours after transfection, cells were transfected with empty vector (EV, pDEST-FRT-FLAG) or I-SceI plasmids. After 48 hours, cells were trypsinized, harvested, washed and re-suspended in PBS. The percentage of GFP-positive cells were determined by flow cytometry. The data was analyzed using the FlowJo software.

### Class Switch Recombination Assay

Immunoglobulin (Ig) M (IgM) to IgA switching was assayed in CH12F3-2 cells with integrated shRNA for REV7, AID, or FAM35A. Cells were activated in 1 mL complete CH12F3-2 media with 1.25 ng transforming growth factor beta 1 (TGF-β1, PeproTech), 5 ng interleukin (IL) 4 (IL-4, PeproTech) and 0.5 μg anti-cluster of differentiation (CD) 40 (CD40, eBioscience). IgA expression was measured by flow-cytometry using primary conjugated anti-mouse IgA-PE (Southern Biotech) at 24h and 48h after activation. Proliferation of the different transduced CH12F3-2 cell lines was monitored using carboxyfluoroscein succinimidyl ester (CFSE, Invitrogen) following the manufacturer’s guidelines. Class switching assays were done in triplicate for every independent experiment.

### Comet assay

U2OS cells were exposed to IR (10 Gy) and processed according to manufacturer’s recommendations (Trevigen). Cells were trypsinized at the indicated time points and resuspended at 10^5^ cells/mL in PBS. Cells were combined with low melting agarose at 1:10 ratio and spread over the CometSlide. Slides were allowed to dry at 4°C for ten minutes then immersed in lysis buffer (Trevigen) overnight. The next day the slides were immersed neutral electrophoresis buffer (two 15-minute washes) followed by electrophoresis at 31V for 45 minutes. Subsequently the slides were incubated for 30 minutes in DNA precipitation solution followed by 30 minutes in 70% ethanol. Slides were dried and stained with SYBR Gold (Invitrogen). Images were taken using the EVOS FL Cell Imaging System microscope and the tail moment was quantified using the CaspLab software. For each condition, at least 50 cells were analyzed.

### Laser Micro-irradiation

U2OS stable cell populations expressing the various constructs were transferred to a 96-well plate with 170 μm glass bottom (Ibidi), presensitized with 10 μg/mL Hoescht 33342 and microirradiated using a FV-3000 Olympus confocal microscope equipped with a 405nm laser line as described previously (Gaudreau-Lapierre et al., 2018). Immunofluorescence was performed as described previously (Gaudreau-Lapierre et al., 2018). Briefly, following microirradiation, cells were allowed to recover before pre-extraction in 1X PBS containing 0.5 % Triton X-100 on ice for five minutes. Following washes with 1X PBS, cells were fixed for 15 min in 3 % paraformaldehyde 2 % sucrose 1X PBS solution, permeabilized in 1X PBS containing 0.5 % Triton X-100 for five min, blocked in 1X PBS containing 3% BSA and 0.05% Tween-20 and stained with the following primary antibodies 1:500 RPA32 mouse (Santa Cruz, sc-56770) or 1:500 γ-H2A.X mouse (abcam, ab26350) and 1:500 HA-tag rabbit (Bethyl, A190-108A). After extensive washing, samples were incubated with 1:250 each of goat anti-mouse Alexa 488-conjugated and goat anti-rabbit Alexa 647-conjugated antibodies (Cell Signaling 4408S and 4414S). DAPI staining was performed and samples were imaged on a FV-3000 Olympus confocal microscope.

### ChIP Quantitive PCR

Stable 293T cell lines expressing ER-AsiSI cells were transfected with Flag-FAM35A and treated with 1 μM of 4-OHT (4-Hydroxytamoxifen) for 6 hours. Cells were collected for ChIP assay as per previously (Iacovoni et al., 2010). Briefly, cells were crosslinked using 1.5 mM EGS (ethylene glycol bis(succinimidyl succinate), Thermo Fisher # 21565), followed by 1 % of formaldehyde. Cell nuclei was isolated and lysed. Chromatin was sonicated for 15 min using a water bath sonicator/bioruptor. Fragmented chromatin bound to FAM35a and γ-H2AX was immunoprecipitated using Anti-FLAG Magnetic Beads (Sigma, M8823), and anti-γ-H2AX (JBW301, EMD-Millipore, Massachusetts, US) in combination with protein A/G magnetic Beads, respectively. Antibody/protein/DNA complexes were then eluted and reverse crosslinked. DNA was purified using QIAquick Kit (Qiagen #28106) and used for qPCR detection with the following oligonucleotides: AsiSI22-distF 5’-CCCATCTCAACCTCCACACT-3’; AsiSI22-distR 5’-CTTGTCCAGATTCGCTGTGA-3’; AsiSI22-ProxF 5’-CCTTCTTTCCCAGTGGTTCA-3’; AsiSI22-ProxR 5’-GTGGTCTGACCCAGAGTGGT-3’. IP efficiency was calculated as percentage of input DNA immunoprecipitated.

### ChIP-sequencing

293T cells transfected with Flag-FAM35A or Flag alone (as negative control) were treated with or without 150 ng/ml of NCS (Sigma) for 5 h, followed by ChIP analysis using anti-FLAG Magnetic Beads, as described previously (see section ChIP Quantitive PCR). Chromatin was sonicated for 40 min to obtain an average length of ~150 bp. Immunoprecipitated DNA was quantified by AccuClear^®^ Ultra High Sensitivity dsDNA Quantitation Kit (Biotium, Inc. #31029) and 100 ng of DNA was used for Next-Generation Sequencing using Illumina HiSeq 2500, paired end sequencing of 125-bases (TCAG Facilities, Toronto). Sequencing data were aligned using BWA and peaks were identified using Macs2. They were further annotated using HOMER.

### DNA Fiber Combing

U2OS cells were transfected with the indicated siRNA in a 6-well cell culture plate. After 48 hours, cells were treated with indicated schedules and concentrations of thymidine analogue pulses (chlorodeoxyuridine (CldU; C6891); iododeoxyuridine (IdU; I7125); Sigma, Missouri, US) with and without neocarzinostatin (NCS, Sigma) treatment to measure replication fork kinetics and extent of DNA end-resection. Cells were trypsinized, agarose plug embedded, and subjected to DNA extraction as per the Fibreprep protocol (Genomic Vision, Bagneux, FR). Vinylsilane coated coverslips (Genomic Vision) were combed through prepared DNA solution using FibreComb Molecular Combing system (Genomic Vision). Combed DNA was dehydrated, denatured, blocked with BlockAid blocking solution (Invitrogen, California, US) and stained with mouse anti-BrdU (B44, BD, New Jersey, US), rat anti-BrdU (BU1/75, Abcam, Cambridge, UK) and rabbit anti-ssDNA (18731, Immuno-Biological Materials, Gunma, Japan) antibodies. Slides were subsequently washed and stained with secondary antibodies: Goat anti-rabbit IgG conjugated Alexa Fluor 480 (BD Horizon) goat anti-mouse IgG conjugated Alexa Fluor 555 (Invitrogen), goat anti-rat conjugated Cy5 (Abcam). Slides were dehydrated, mounted, and visualized using FibreScan services (Genomic Vision).

### Phospho-H2AX Flow Cytometry

U2OS cells were transfected with indicated siRNA in a 6-well cell culture plate. After 48 hours, cells were treated with NCS for 30 mins and after indicated time intervals, cells were trypsinized, washed, and fixed with 1% para-formaldehyde, washed and subsequently permeabilized in 70% ethanol at −20°C. Cells were washed twice with intracellular wash buffer (1% BSA, 0.05% Tween-20, PBS) and re-suspended in 1.0 ug/mL mouse anti-γ-H2AX (JBW301, EMD-Millipore, Massachusetts, US) for one hour at RT. Cells were then washed and re-suspended in 2.0 ug/mL goat anti-mouse Alexa Fluor 647 (Invitrogen) for one hour at RT. Cells were washed and re-suspended in a propidium iodide (PI) solution (20 ug/mL PI, 300ug/mL RNase, PBS), incubated at RT for 30 minutes. Events were acquired on a LSRFortessa (BD). Events were analyzed on FlowJo v10 (Treestar, Oregon, US).

### CRISPR/Cas9 genome-wide screen

For the genome-wide CRISPR/Cas9-based screen, 270 million RPE-hTERT/Cas9 cells were transduced as described previously (Hart et al., 2015) with TKOv1 concentrated library virus at MOI = 0.2, ensuring a coverage of at least 600-fold for each individual sgRNA represented in the cell population. Two days later, puromycin was added to the media at a final concentration of 15 ug/ml and incubated for four days to allow for the emergence of resistant cells with fully repaired sgRNA library targeted loci. Cells were then split into 2 pools each in triplicate at a cell density of 54 million cells/replicate and treated with either vehicle (H_2_0) or doxorubicin at its IC25 (3 nM) and cultured for two weeks with puromycin at a concentration of 7.5 ug/mL. Cells were passaged every three days keeping a minimum cell concentration of 54 million cells per replicate to ensure that a 600-fold library coverage was maintained over the duration of selection. At each time point, cell pellets were collected and frozen prior to genomic DNA extraction. Cell pellets were resuspended in 6 mL DNA lysis buffer (10 mM Tris-Cl, 10mM EDTA, 0.5% SDS, pH 8.0) with 100 ug/mL RNase A, followed by incubation at 37 °C for 60 min. Proteinase K was subsequently added (400 ug/mL final) and lysates were further incubated at 55 °C for two hours. Samples were then briefly homogenized by passing them three times through a 18G needle followed by three times through a 22G needle. Sheared samples were transferred into pre-spun 15 mL MaXtract tubes (Qiagen) mixed with an equal volume of neutral phenol:chlorophorm:isoamyl alcohol (25:24:1) solution, shook and centrifuged at 1,500g for five min at RT. The aqueous phase was extracted and precipitated with two volumes of ethanol and 0.2M NaCl. Air-dried pellets were resuspended in water and quantitated via UV absorbance spectrometry.

For next-generation sequencing (NGS), sgRNA integrated loci were amplified from 330 ug of total genomic DNA per replicate using two rounds of nested PCR. The initial outer PCR consisted of 25 cycles with an annealing temperature of 65 °C using Hot start Q5 polymerase (NEB) using primers Outer Primer Forward (AGGGCCTATTTCCCATGATTCCTT) and Outer Primer Reverse (TCAAAAAAGCACCGACTCGG). PCR reactions were pooled and ~2% of the input was amplified a further 12 cycles for the addition of Illumina HiSeq adapter sequences. The resulting ~200bp product from each pooled sample was further purified following separation in a 6% 0.5XTBE polyacrylamide gel. The amplicon library NGS-ready final product was quantified using qPCR and submitted for deep-sequencing on the HiSeq 2500 Illumina platform using standard Single-Read (SR) 50-cycle chemistry with dual-indexing with Rapid Run reagents. The first 20 cycles of sequencing were "dark cycles", or base additions without imaging. The actual 26bp read begins after the dark cycles and contains two index reads, reading the i7 first, followed by i5 sequences. Prior to analysis, FastQ NGS read files were initially processed using FastQC software to assess uniformity and quality. Reads were trimmed of NGS adapter sequences using the Cutadapt tool. Reads were aligned to the sgRNA library index file using Bowtie to assign a matching gene-specific sgRNA, and total read count tables were subsequently generated using Samtools. A pseudocount of 1 was added to each sgRNA read count, and reads were normalized to the total read count per experimental replicate. Any sgRNA that had fewer than 25 total reads in any replicate or which were represented by less than 3 unique sgRNA for a given gene were dropped from the analysis. Average log2 fold-change was calculated for a given gene between the initial and final abundances for all sgRNAs targeting it across the replicates.

### Phylogenetic profiling analysis

To identify genes co-evolved with FAM35A we used normalized phylogenetic profiling as previously described (Tabach et al., 2013a; Tabach et al., 2013b). Briefly, we have generated the phylogenetic profile of 42 mammalian species and calculated the Pearson correlation coefficients between the phylogenetic profile of FAM35A and the phylogenetic profiles of 19520 human protein coding genes, and defined the 200 genes with the highest correlation coefficients as co-evolved with FAM35A in mammalians. In a similar manner, we identified the top 200 genes that co-evolved with FAM35A in 63 vertebrate species. The intersection between these two lists yielded 159 genes that were subsequently considered as co-evolving with FAM35A with high confidence and further processed for pathway enrichment analysis.

### Patient Cohort analysis

TNBC patient data and PDXs were collected in accordance with the McGill University Health Center research ethics board (SUR-99-780) and the McGill University Animal Care Committee (2014-7514) guidelines. The Mean-Centered Expression as well as a survival data was included 30 primary tumors and 30 pdx samples. For the following analyses we have included expression from only the 24 primary tumor samples. Six of the patients were removed from the analyses. One patient died on the day of surgery and another 5 samples had no follow-up time. Survival analysis was performed using the coxph function in the R package ‘survival’. Expression was classified as either low or high using the top quartile as the threshold.

## ACKNOWLEDGMENTS

We are grateful to Amelie Fradet-Turcotte, Michael Witcher, Josie Ursini-Siegel, Chantal Autexier and William Foulkes for critical reading of the manuscript; to Daniel Durocher, Anne-Claude Gingras, Jeremy Stark, Michael Witcher and Roger Greenberg for plasmids and other reagents. We would like to specifically thank Roderick McInnes, Josie Ursini-Siegel and Koren Mann for their constant support. JH, VL and MK received a doctoral fellowship from the Cole Foundation. AM was supported by a post-doctoral fellowship from the Cole Foundation. ESC received a FRQS postdoctoral training scholarship. HB was supported by a doctoral training award from the FRQS (#33603). JFC is the recipient of the TRANSAT chair in Breast Cancer Research. AO is the Canada Research Chair (Tier 2) in Genome Stability and Hematological Malignancies. Work in the AO laboratory was supported by a CIHR Project Grant (#376245), a CRS Operating Grant (#21038), a Transition Grant from the Cole Foundation and an internal Operating Fund from the Sir Mortimer B. Davis Foundation of the Jewish General Hospital. Work in the JFC laboratory was supported by a NSERC Discovery Grant (RGPIN-2016-04808). Work in the AM laboratory was supported by a NSERC Discovery Grant (#5026) and a CIHR Project Grant (#376288). “Life always offers you a second chance. It is called tomorrow!”

## AUTHOR CONTRIBUTIONS

SF and JH designed, performed most of the experiments presented in this manuscript and analyzed the data. VL designed and performed the CSR experiments and analyzed the data. AB designed, performed the CRISPR/Cas9-based genome-wide screen and the clonogenic experiments and analyzed the data. TM and BD performed the laser stripe micro-irradiation and AM designed the experiments and analyzed the data. ZL generated most of the constructs used in this manuscript, designed and performed the ChIP experiments and analyzed the data. AS designed and performed the comet assay experiments and analyzed the data. ESC, HB, DG, CD and HK performed the MS experiments and analyzed the data under the supervision of JFC. DR performed the phylogenetic analysis profiling under the supervision of YT. KM performed the pathway analysis of FAM35A. KK performed the RNA-seq analysis and the patient cohort analysis under the supervision of CMG. MP provided the RNA-seq data and the related patient outcome of the TNBC cohort. AO conceived the study, designed the research, provided supervision and wrote the manuscript with input from all the other authors.

## SUPPLEMENTARY FIGURE LEGENDS

**Supplemental Figure 1.**
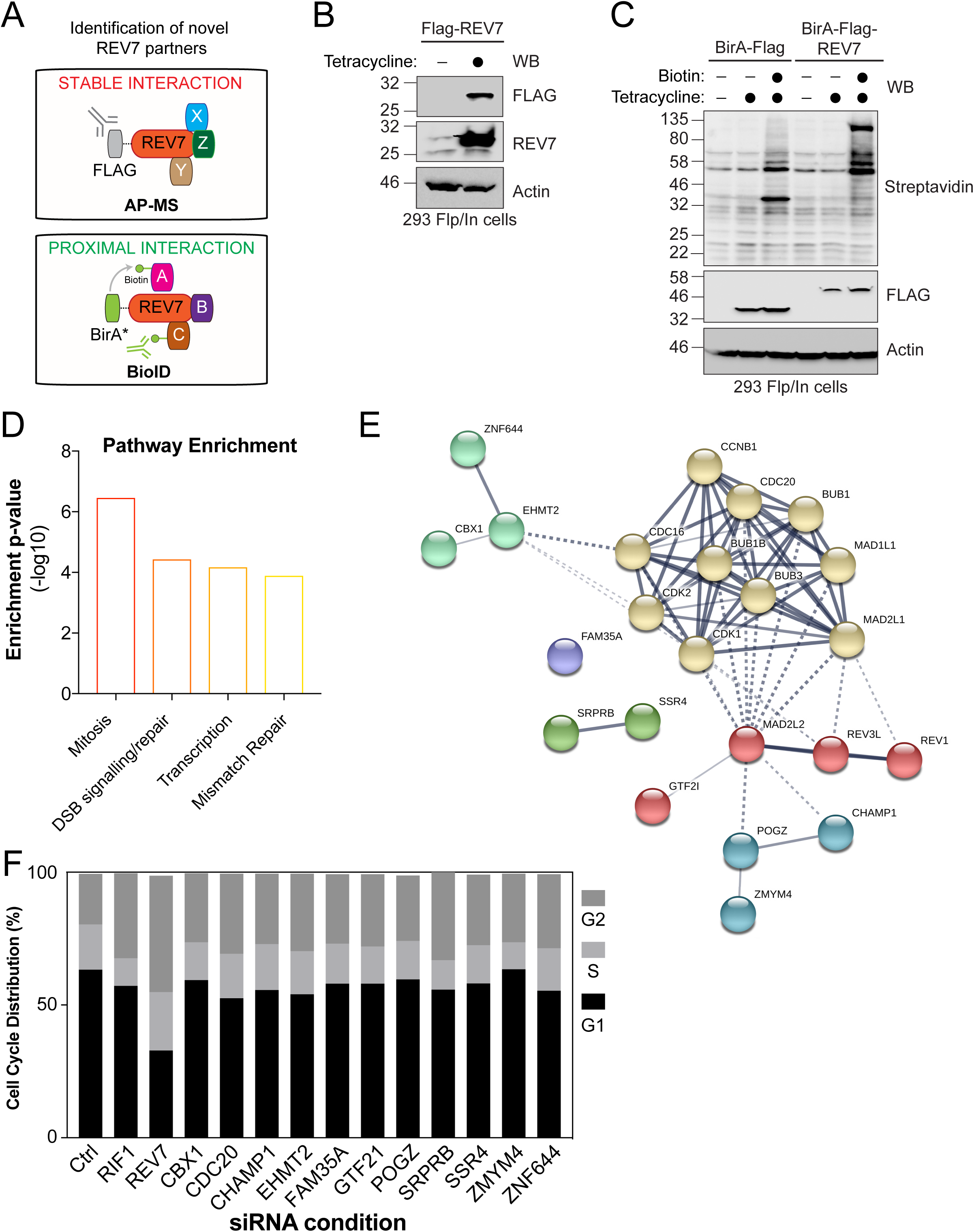
(A) Schematic representation of AP-MS stable interaction Flag-REV7 pulldown and the BioID BirA*-REV7 biotinylation of proximal interactors. (B) HEK293-TREx cells stably expressing an inducible Flag-REV7 construct were tested for expression following induction with tetracycline as indicated. After lysis, samples were immunoblotted for FLAG and REV7. Actin was used as a loading control. (C) HEK293-TREx cells stably expressing an inducible BirA-Flag or BirA-Flag-REV7 construct were tested for expression and biotinylation following induction with tetracycline and incubation with biotin as indicated. After lysis, samples were immunoblotted for FLAG and Streptavidin. Actin was used as a loading control. (D) The interactome of REV7 obtained from both the AP-MS and the BioID approaches were analyzed for pathway enrichment using EnrichR. The y-axis represents the ratio of the number of genes from the dataset that map to the pathway and the number of all known genes ascribed to the pathway and is defined as enrichment of p-value(-log10). (E) Network representation of the selected 11 high-confidence interactors of REV7 (annotated as MAD2L2 in this figure) and their previously described interactors. Proteins are represented following the k-means clustering through STRING v10.5. (F) Cell cycle distribution of U2OS EJ5-GFP cells transfected with the indicated siRNAs and subsequently for Propidium iodide (PI) staining and flow cytometry analysis. Data are presented as the mean (n=2).

**Supplemental Figure 2.**
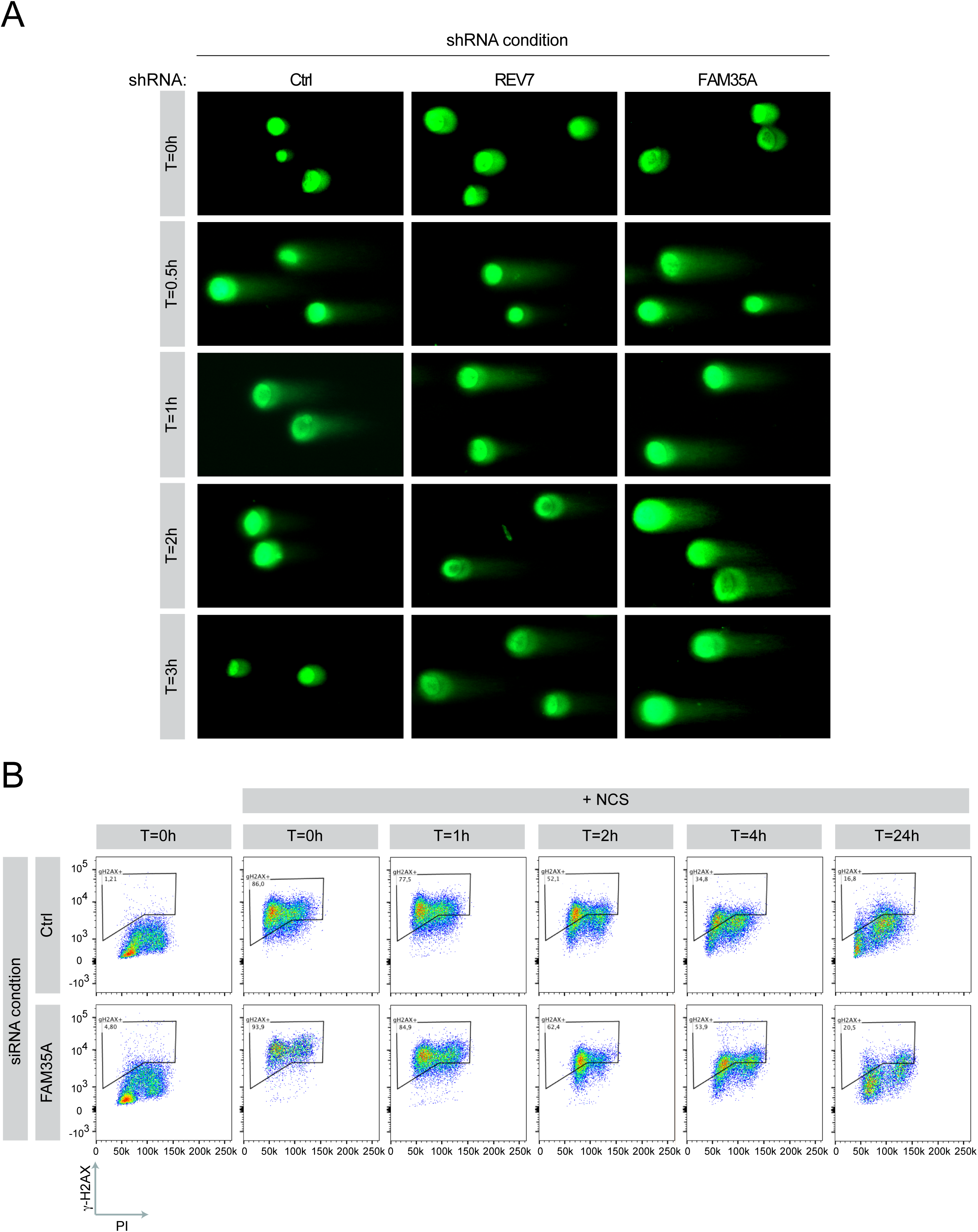
(A) Representative images of Comet Assay Tail Moment quantified in Figure 2. U2OS cells stably expressing shCtrl, shREV7 or shFAM35A were exposed to irradiation (10Gy) and run in low melting agarose under neutral conditions. Immunofluorescence against DNA stained with SYBR Gold was performed to measure the tail moment. (B) Representative flow cytometry profiles of U2OS cells transfected with the indicated siRNA and subsequently treated with NCS for 30 mins before being trypsinized and processed for anti-γ-H2AX (y axis) and PI (x axis) staining.

**Supplemental Figure 3.**
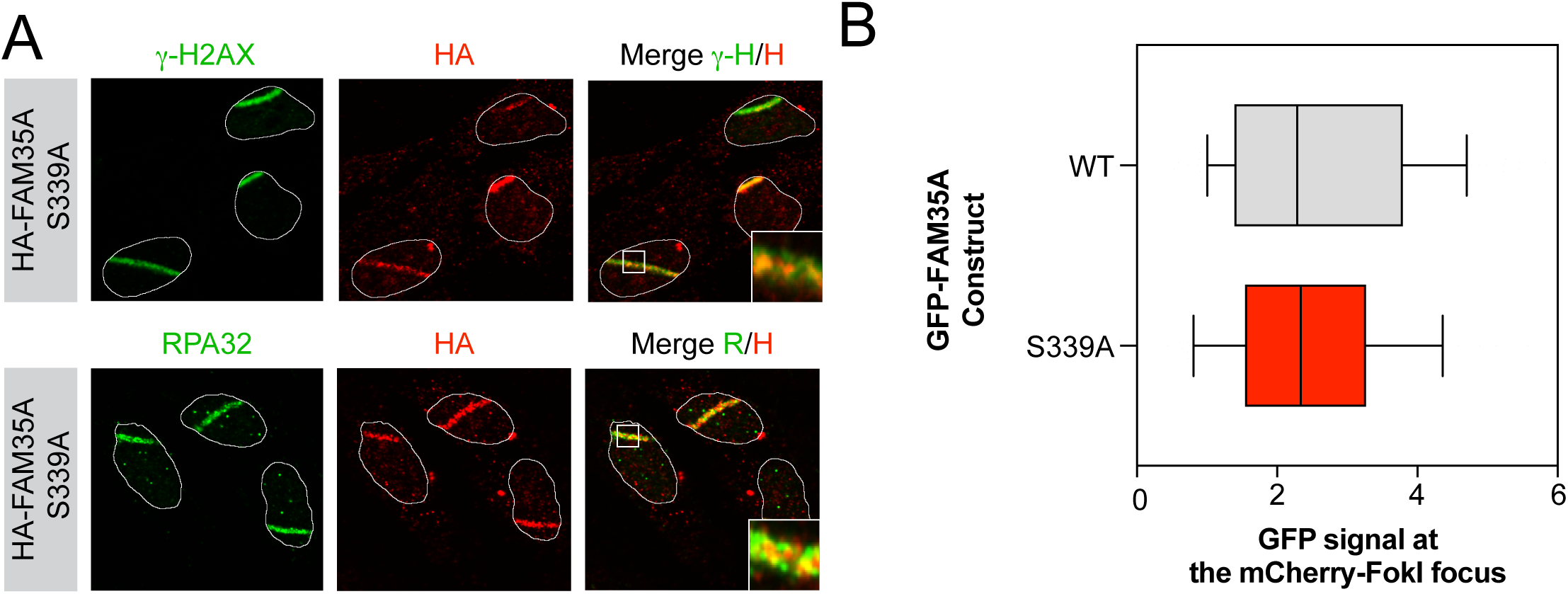
(A) U2OS cells stably expressing HA-FAM35A-S399A were processed as previously described Immunofluorescence against endogenous HA, γ-H2AX (Top) and RPA32 (Bottom) epitope was subsequently performed to monitor their accumulation at sites of damage. Shown are representative micrographs. (B) U2OS LacR-Fok1 cells were transfected with GFP-FAM35A or GFP-FAM35A-S339A mutant and 24 h later DNA damage was induced using Shield-1 and 4-OHT. The cells were then processed for GFP and mCherry immunofluorescence. Shown is the quantification of cells expression GFP at Fok1 sites. Data are represented as a box-and-whisker plot where the whiskers represent the 10-90 percentile. At least 75 cells were counted per condition. Significance was determined by one-way ANOVA followed by a Dunnett test.

**Supplemental Figure 4.**
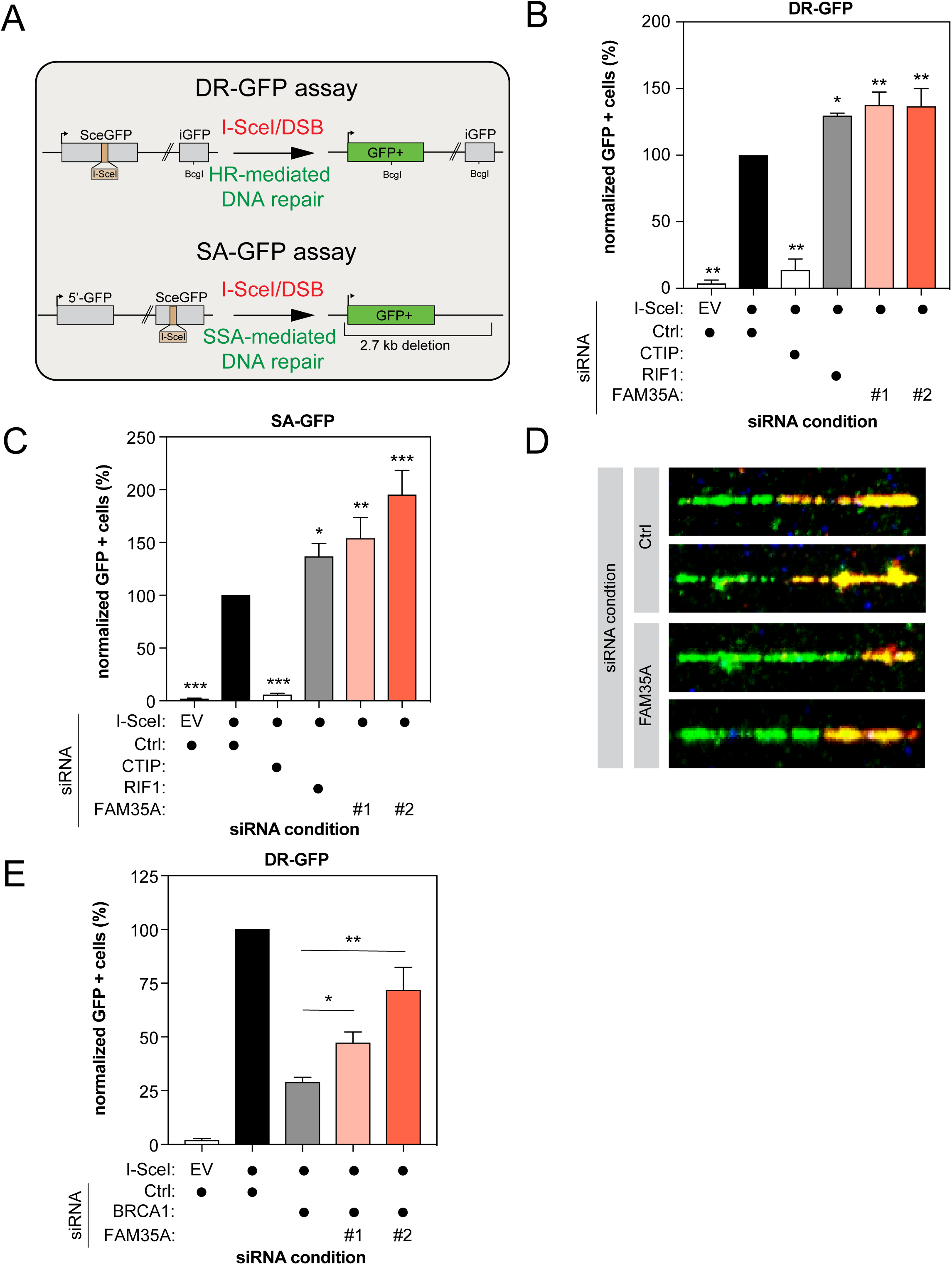
(A) Schematic diagram of both the DR-GFP reporter assay (Top) and the SA-GFP reporter assay showing (Bottom). (B) HeLa DR-GFP cells were transfected with the indicated siRNAs. At 24hr post-transfection, cells were transfected with the I-SceI expression plasmid, and the GFP^+^ population was analyzed 48 hr post-plasmid transfection. The percentage of GFP^+^ cells was determined for each individual condition and subsequently normalized to the non-targeting condition (siCTRL). Data is presented as the mean ± SD (n=3). Significance was determined by one-way ANOVA followed by a Dunnett test. *p<0.0005, **p=0.0001. (C) U2OS SA-GFP cells were transfected with the indicated siRNAs. At 24hr post-transfection, cells were transfected with the I-SceI expression plasmid, and the GFP^+^ population was analyzed 48 hr post-plasmid transfection. The percentage of GFP^+^ cells was determined for each individual condition and subsequently normalized to the non-targeting condition (siCTRL). Data is presented as the mean ± SD (n=3). Significance was determined by one-way ANOVA followed by a Dunnett test. *p<0.05, **p<0.005, ***p<0.0005. (D) Representative images of the DNA fiber assay obtained from U2OS cells which were transfected with the indicated siRNAs and then treated with CldU, IdU and NCS 48h post-transfection as indicated. The slides were stained, dehydrated, mounted and visualized and shown is the quantification of CldU/IdU tract length in order to visualize DNA end-resection. (E) HeLa EJ5-GFP cells were co-transfected with siBRCA1 and the indicated siRNAs. At 24hr post-transfection, cells were transfected with the I-SceI expression plasmid, and the GFP^+^ population was analyzed 48 hr post-plasmid transfection. The percentage of GFP^+^ cells was determined for each individual condition and subsequently normalized to the non-targeting condition (siCTRL). Data is presented as the mean ± SD (n=3). Significance was determined by one-way ANOVA followed by a Dunnett test. *p<0.05, **p<0.0005.

**Supplemental Figure 5.**
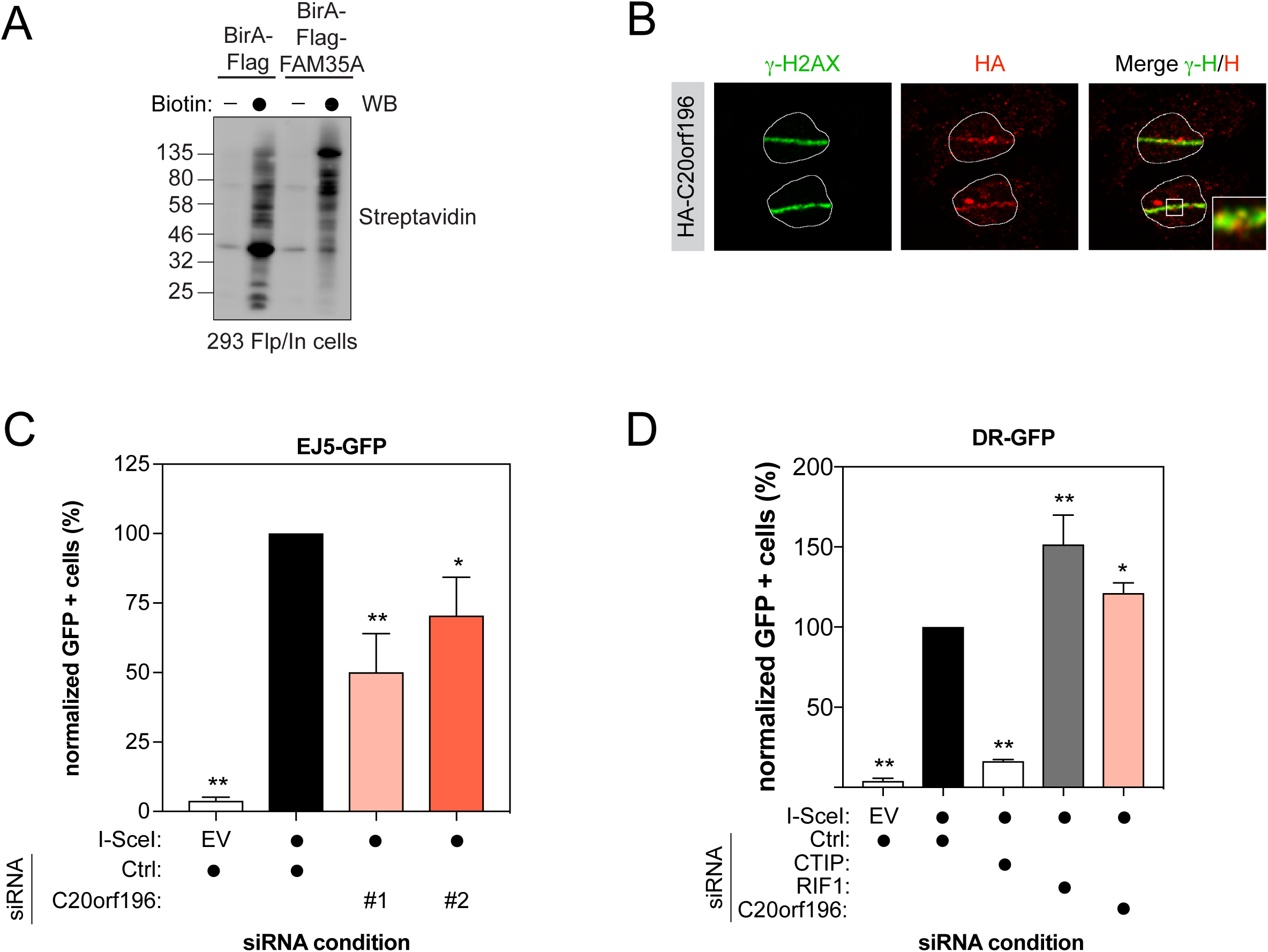
(A) HEK293-TREx cells transfected with a BirA-Flag or BirA-Flag-FAM35A construct were tested for biotinylation following incubation with biotin as indicated. After lysis, samples were immunoblotted for Streptavidin. (B) U2OS cells stably expressing HA-C20orf196 were processed as previously described Immunofluorescence against endogenous HA and γ-H2AX epitope was subsequently performed to monitor their accumulation at sites of damage. Shown are representative micrographs. (C) U2OS EJ5-GFP cells were transfected with the indicated siRNAs. At 24hr post-transfection, cells were transfected with the I-SceI expression plasmid, and the GFP^+^ population was analyzed 48 hr post-plasmid transfection. The percentage of GFP^+^ cells was determined for each individual condition and subsequently normalized to the non-targeting condition (siCTRL). Data is presented as the mean ± SD (n=3). Significance was determined by one-way ANOVA followed by a Dunnett test. *p<0.005, **p<0.0005. (D) U20S DR-GFP cells were transfected with the indicated siRNAs. At 24hr post-transfection, cells were transfected with the I-SceI expression plasmid, and the GFP^+^ population was analyzed 48 hr post-plasmid transfection. The percentage of GFP^+^ cells was determined for each individual condition and subsequently normalized to the non-targeting condition (siCTRL). Data is presented as the mean ± SD (n=3). Significance was determined by one-way ANOVA followed by a Dunnett test. *p<0.05, **p<0.0005.

